# Scalable computation of ultrabubbles in pangenomes by orienting bidirected graphs

**DOI:** 10.64898/2026.03.28.714704

**Authors:** Juha Harviainen, Francisco Sena, Corentin Moumard, Aleksandr Politov, Sebastian Schmidt, Alexandru I. Tomescu

## Abstract

**Motivation:** Pangenome graphs are increasingly used in bioinformatics, ranging from environmental surveillance and crop improvement to the construction of population-scale human pangenomes. As these graphs grow in size, methods that scale efficiently become essential. A central task in pangenome analysis is the discovery of variation structures. In directed graphs, the most widely studied such structures, *superbubbles*, can be identified in linear time. Their canonical generalization to bidirected graphs, *ultrabubbles*, more accurately models DNA reverse complementarity. However, existing ultrabubble algorithms are quadratic in the worst case.

**Results:** We show that all ultrabubbles in a bidirected graph containing at least one tip or one cutvertex—a common property of pangenome graphs—can be computed in linear time. Our key contribution is a new linear-time orientation algorithm that transforms such a bidirected graph into a directed graph of the same size, in practice. Orientation conflicts are resolved by introducing auxiliary source or sink vertices. We prove that ultrabubbles in the original bidirected graph correspond to weak superbubbles in the resulting directed graph, enabling the use of existing linear-time algorithms. Our approach achieves speedups of up to 25×over the ultrabubble implementation in `vg`, and of more than 200× over `BubbleGun`, enabling scalable pangenome analyses. For example, on the v2.0 pangenome graph constructed by the Human Pangenome Reference Consortium from 232 individuals, after reading the input, our method completes in under 3 minutes, while `vg` requires more than one hour, and four times more RAM.

**Availability:** Our method is implemented in the BubbleFinder tool github.com/algbio/BubbleFinder, via the new ultrabubbles subcommand.

## 1 Introduction

### 1.1 Background

Genomic variations induce specific subgraph structures in genome graphs. For instance, in de Bruijn graphs, single nucleotide polymorphisms typically induce *bubbles* (Zerbino and Birney, 2008; Fasulo et al., 2002) consisting of two paths of the same length sharing the same start and end vertices. In genome assembly, the most prominent generalization of bubbles are *superbubbles* (Onodera et al., 2013). These are defined as acyclic subgraphs of a directed graph with a unique entry vertex from which all internal vertices are reachable, and a unique exit vertex reachable from all internal vertices, such that all edges connecting the super-bubble to the rest of the graph are incident to the entry or exit vertices. Superbubbles have numerous applications, see e.g. (Kolmogorov et al., 2020; Rautiainen et al., 2023; Minkin and Medvedev, 2020).

The development of the pangenomic field has led to renewed interest in defining and efficiently computing graph structures capturing genomic variation. A particular focus has also been on directly defining these structures in *bidirected* graphs (or in equivalent structures, such as *biedged graphs*), which innately capture the reverse complementarity of the DNA. In a bidirected graph, bidirected edges have a sign (+ or−) at each endpoint, and e.g. bidirected walks must alternate sign when passing through a vertex.

The “canonical” generalization of superbubbles to bidirected graphs are *ultrabubbles*, introduced by Paten et al. (2018). They are analogously defined as subgraphs connected to the rest of the graph by two designated endpoints. Within an ultrabubble, all edges incident at each endpoint must have the same sign internal to the bubble, while edges of the opposite sign connect to the rest of the graph. Furthermore, ultrabubbles are internally acyclic (containing no bidirected walks that start and end at the same vertex) and contain no internal tips (vertices whose incident edges all have the same sign).

### 1.2 Motivation

A significant drawback of ultrabubbles compared to superbubbles is that the currently best algorithm computing all of them takes *O*((|*V* | + |*E* |)^2^) time, in the worst case, for a bidirected graph (*V, E*) (Paten et al., 2018). An implementation finding all ultrabubbles exists in the `vg` toolkit (Garrison et al., 2018), based on first finding a set of *snarls* (a superset of ultrabubbles) of size linear in the size of the graph (a *snarl decomposition*), and then checking which of these snarls is an ultrabubble.

Interestingly, we note that (weak) superbubbles can also be applied to bidirected graphs by considering their “doubled” directed graph: every vertex-side *v*+ and *v*− is modeled by a separate vertex, and directed edges are added to and from *v*+ or *v*− based on the signs of each bidirected edge incident to *v* (see the Supplementary Material for details). Conceptually, the Python-based `BubbleGun` tool by Dabbaghie et al. (2022) computes, and merges, pairs of matching (weak) superbubbles in this doubled graph. While in our experiments we observed that the number of resulting bubbles matches the number of ultrabubbles, to the best of our knowledge we are unaware of a proof that this would lead to an algorithm for computing *all* ultrabubbles (in fact, such proof would immediately imply that ultrabubbles can be computed in linear time). However, an innate drawback of this approach is that it works on a graph of double the size, leading to double the memory and the time needed to compute superbubbles in it (slowdown which we also confirm experimentally).

Recently, other types of bubble-like structures have been proposed, for example *bibubbles*, introduced by Li et al. (2024) and *panbubbles*, introduced by Bhat et al. (2025), none of which admit linear-time algorithms computing all such structures. These have been shown useful in capturing biological events in complex gene graphs. On the other hand, *snarls*, also introduced by Paten et al. (2018), have been recently shown to admit a linear-time algorithm identifying them all, via SQPR trees (Sena et al., 2025).

As pangenomes grow, scalability becomes paramount, making linear-time algorithms essential. For example, Release 1 (May 2023) of the Human Pangenome Reference Consortium (HPRC) graph contained 47 individuals (94 haplotypes) and 110 million edges (Liao et al., 2023). Release 2 (May 2025) expanded to 232 individuals^1^ and 206 million edges, while Release 3, scheduled for Summer 2026, is expected to include over 350 individuals.^2^

### 1.3 Results

In this paper, we make progress in the direction of obtaining linear-time algorithms for bubble-like structures, if we assume that the pangenome graph has at least one tip (a vertex whose incident bidirected edges all have the same sign), or at least one cutvertex. In such a graph, we prove that all ultrabubbles can be computed in linear time *O*(|*V* | +|*E*|), significantly improving over the existing *O*((| *V*| + |*E*|)^2^)-time algorithm by Paten et al. (2018). This improves also over the recent method by Zisis and Sætrom (2026) of verifying which snarls in a given set are ultrabubbles, running in time *O*(*K* |*V*| + (|*V* |+ |*E*|)), where *K* is the number of given snarls. While the existence of a tip or a cutvertex is limiting in theory, pangenome graphs are rarely tipless, as also mentioned by Bhat et al. (2025), and as we also observed in our experiments, where all graphs had either a tip or a cutvertex.

We obtain this result by showing that computing ultrabubbles can be reduced, in linear time, to computing *weak* superbubbles, a type of bubble which functions as a slight relaxation of superbubbles. Our approach relies on a novel algorithm that transforms a bidirected graph into an equivalent directed graph. Starting from a tip or a cutvertex, the algorithm performs a depth-first search that *orients* the bidirected graph. Orientation is achieved by *flipping* the signs incident to some vertices so that every edge has opposite signs at its endpoints; such edges can then be interpreted as directed edges.

We show that ultrabubbles admit such an orientation, and that once oriented they correspond to weak superbubbles in the resulting directed graph. In cases where orientation conflicts arise—namely, edges whose endpoints have the same sign and for which neither endpoint can be flipped— we resolve these conflicts by replacing the problematic edges with edges incident to newly introduced tips. These new tips correspond to sources or sinks in the directed graph. Although sources and sinks are not inside weak superbubbles by definition, the presence of such a conflict implies that the original structure could not have been an ultrabubble. We further show that the size of the resulting directed graph remains linear in the size of the original bidirected graph (and having only at most 0.2% additional vertices in the HPRC graphs), that ultrabubbles in the bidirected graph correspond to weak superbubbles in the directed graph, and that essentially all weak superbubbles in the directed graph correspond to ultrabubbles in the bidirected graph. Since ultrabubbles are the canonical generalization of superbubbles (Paten et al., 2018), such a relationship must have existed in principle, although it had not previously been made explicit.

Our orientation algorithm is also very simple and efficient. We implemented it into our `BubbleFinder` tool (Sena et al., 2025) (via the new subcommand `ultrabubbles`), and we used the direct linear-time weak superbubble-finding algorithm by Gärtner and Stadler (2019), via its existing C++ implementation. On the HPRC Release 1 and Release 2 pangenome graphs in GBZ format, after reading the input (with the same parsing library), `BubbleFinder` is 25× faster than `vg`, while using four times less RAM (i.e., 14GiB for Release 1 and 24.8 GiB for Release 2 graphs). On the HPRC Release 1 graph in GFA format, `BubbleFinder` is 200× faster than `BubbleGun`, using three times less RAM.

## 2 Methods

### 2.1 Preliminaries

A *bidirected graph G* = (*V, E*) has vertex set *V* = *V* (*G*) and bidirected edge set *E* = *E*(*G*). A *sign* is a symbol *α* ∈ {+, −}, and the *opposite sign*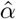of *α* is defined as 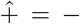and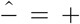. A pair (*v, α*) where *v* ∈ *V* (*G*) and *α* ∈ {+, −}, is a *vertex-side* (Rahman and Medvedev, 2022) (this is also called *signed vertex* in (Abrishami et al., 2025)), which we concisely write as *vα*, e.g. *v*+ or *v*−. A *bidirected edge e ∈ E*(*G*) is an unordered pair of vertex-sides *uα, vβ* ; for simplicity we may refer to bidirected edges as just edges. We say that *G* has a vertex-side of sign *α* in *u* or that *u* has sign *α* over *e*. The set of vertex-sides of *G* is the set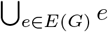. We let 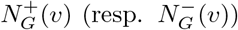denote the set of those vertices *u* for which there is an edge {*v*+, *uα*} (resp. {*v*−, *uα*}) in *G*. We say that a bidirected graph *H* is a *subgraph* of *G* (and write *H* ⊆*G*) if *V* (*H*) ⊆ *V* (*G*) and *E*(*H*) ⊆ *E*(*G*) (we also say that *G* is a *supergraph* of *H*). We say that *v* is a *tip* in *G* if no two vertex-sides of *G* have distinct signs in *v*; a vertex *v* is a *source* (resp. *sink*) if all the vertex-sides of *G* have sign + (resp. —) in *v*. The subgraph *induced* by a subset *C* ⊆ *V* (*G*) of vertices of *G* is the graph with vertex set *C* and the subset of edges in *E*(*G*) whose endpoints are in *C*, which is denoted by *G*[*C*].

A walk *W* in *G* is a sequence of edges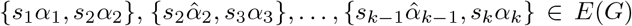. We say that *W* is an *s*_1_*α*_1_*-s*_k_*α*_k_ *walk* or just an *s*_1_*-s*_k_ walk. Vertices *s*_1_ and *s*_k_ are the *first* and *last* vertices of the walk, respectively, and all remaining vertices are its *internal* vertices. Notice that to any walk *W* in a bidirected graph we can associate another walk consisting of the sequence of edges of *W* written in reversed order (i.e., walks in bidirected graphs have two possible orientations). A *path* is a walk without repeated vertices. A *cycloid* is a path where the first and last vertex coincide and where at most one vertex (informally, the *exceptional* vertex) has the same sign over the two edges incident to it in the sequence (see Figure 1). A graph with no cycloid is *acyclic*. A vertex *w* is a *u-v cutvertex* in a bidirected graph *G* if every path between *u* and *v* in the underlying undirected graph of *G*, i.e., if every *undirected path* between *u* and *v*, contains *w* and *w* ≠*u, v*. Graph *G* is *bi-connected* if for any two vertices *u, v ∈ V* (*G*) there is no *u*-*v* cutvertex (such a vertex can also be simply called a *cutvertex* of *G*). Directed graphs can be seen as a special case of bidirected graphs where every edge {*uα, v β* }has *α*≠*β*. More precisely, a *directed graph D* has vertex set *V* (*D*) and *directed edges* in *V* ×*V*. A directed edge from *u* to *v* is denoted as *uv* or (*u, v*), and a *directed path* from *u*_1_ to *u*_k_ in *D* is a sequence of edges *u*_1_*u*_2_, *u*_2_*u*_3_, …, *u*_k 1_*u*_k_ ∈*E*(*D*).

**Figure 1.**
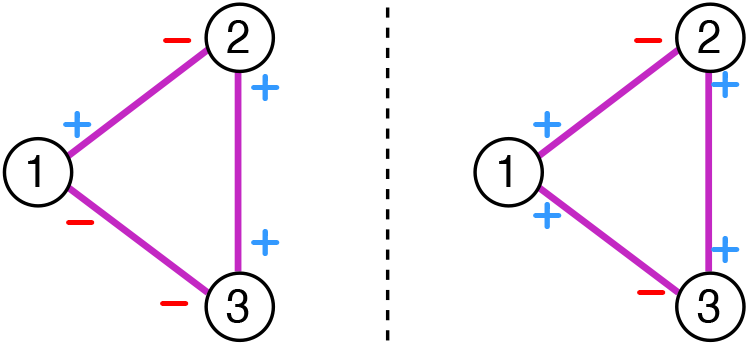
Examples of cycloids in bidirected graphs. On the left, the cycloid {1+, 2−}, {2+, 3+},{3−, 1−} where the sign alternates at every vertex. On the right, the cycloid {1+, 2− }, {2+, 3+}, {3−, 1+}where vertex 1 is the only vertex where the sign does not alternate (i.e., the exceptional vertex). In this paper we say that a bidirected graph is acyclic if it has no cycloid.

Let *v* ∈*V* (*G*) be a vertex. If *vα*_1_, …, *vα*_k_ are all the vertex-sides of *G* containing *v*, then *flipping v* in *G* amounts to replacing each vertex-side *vα*_i_ with 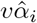 for each *i* ∈ {1, …, *k* }; importantly, the identity of the edges of *G* does not change during this process. A bidirected graph is called *digraphic* (Kita, 2017) if there is a sequence of flips such that each edge of the resulting graph, say *G*^′^, has vertex-sides of opposite signs. If *D* is the directed graph with the same vertex set as *G* (and *G*^′^) and with edge set ∈ {*uv* : {*u*+, *v*−} *E* ∈ (*G*^′^) }, then we say that *D* is an *orientation* of *G*.

Without loss of generality we assume that the underlying undirected graph of our bidirected graphs consist of a single component, since in general we can independently apply our techniques to each component of an arbitrary bidirected graph.

#### Definition 1

(Superbubbloid and Superbubbloid component (Gärtner et al., 2018; Gärtner and Stadler, 2019)). *Let G be a directed graph. Let* (*u, v*) *be an ordered pair of distinct vertices of G. Then* (*u, v*) *is a* superbubbloid *with* entry *u and* exit *v if:*

a. reachability: *G has a directed path from u to v*.
b. matching: *the set of vertices reachable from u without using v as an internal vertex coincides with the set of vertices reaching v without using u as an internal vertex. The graph B is the induced subgraph of G by this set of vertices and it is called the* superbubbloid component.
c. acyclicity: *B is acyclic*.

We also need the following notions introduced in (Gärtner et al., 2018; Gärtner and Stadler, 2019). A *weak superbubbloid* is a pair (*u, v*) with *u* ≠ *v* such that (*u, v*) satisfies conditions (a) and (b) of superbubbloid (say, with graph *B* from condition (b)), and such that *B*− *vu* is acyclic. A *(weak) superbubble* is a (weak) superbubbloid that is minimal in the sense that there is no vertex *w* ∈ *V* (*B*) distinct from *v* such that (*u, w*) is a (weak) superbubbloid (see Figure 2a). A *trivial (weak) superbubble* (*u, v*) is a (weak) superbubble whose vertex set is *u, v*. For clarity, notice that any superbubbloid is also a weak superbubbloid since *vu* may not be an edge.

**Figure 2.**
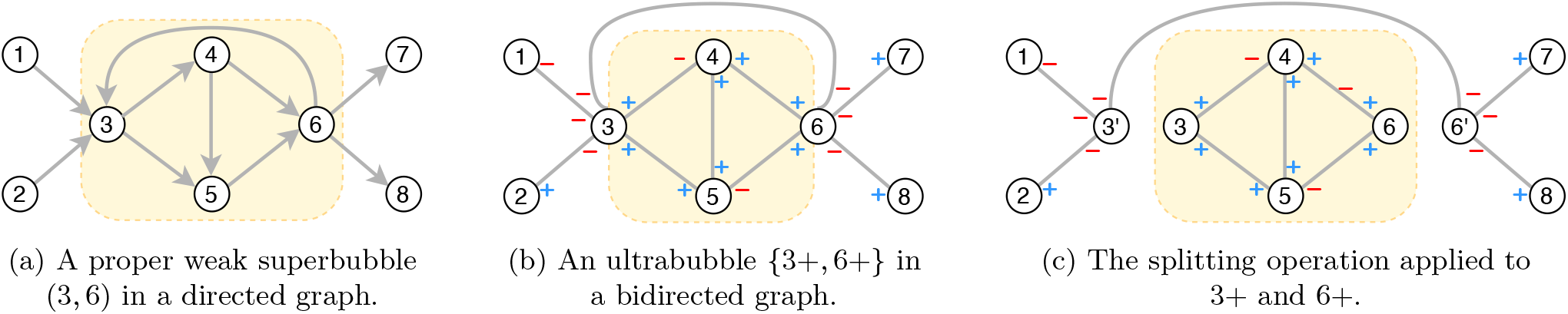
Examples of (weak) superbubbles and ultrabubbles. (a): A proper weak superbubble (3, 6) with component having vertex set {3, 4, 5, 6}, in yellow. Without the edge (6, 3), the weak superbubble (3, 6) becomes a superbubble. (b): An ultrabubble {3+, 6+ }with ultrabubble component having vertex set{3, 4, 5, 6,} in yellow. Note that the signs at 3 and 6 are partitioned according to the ultrabubble component. (c): Applying the splitting operation. For 3+ we add a new node 3^′^ such that all +-incident edges remain at 3 and all−-incident edges move to 3^′^ (and analogously for 6+).

To define ultrabubble we need one more definition. The *splitting* operation receives a bidirected graph *G* and a vertex-side *uα* and produces a new bidirected graph *G*^′^ = (*V* ^′^, *E*^′^) with *V* ^′^ := *V* (*G*) ∪ {*u*^′^}and *E*^′^ := *E*(*G*) \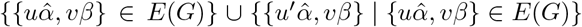. Essentially, every edge of *G* incident to *u* with sign 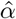will be incident to *u*^′^ in *G*^′^ instead, and those edges incident to *u* with sign *α* remain unchanged from *G* to *G*^′^ (see Figure 2c), so *u* and *u*^′^ become tips of opposite signs in *G*^′^.

Ultrabubbles were defined by Paten et al. (2018) in “biedged graphs”. In fact, ultrabubbles are “snarls” (see (Paten et al., 2018)) whose component is acyclic and whose “interior” contains no tips. Originally, Paten et al. (2018) defines acyclicity as the absence of closed walks where at most one vertex is allowed to be entered and left with the same sign. Here, for simplicity, we define acyclicity as the absence of cycloids, and is not hard to see that these two notions are equivalent.^3^ Further, here we use an equivalent definition of ultrabubble for bidirected graphs based on the definition of “snarl” given in (Sena et al., 2025).

#### Definition 2

(Ultrabubble and Ultrabubble component (Paten et al., 2018; Sena et al., 2025), see Figure 2b). *Let G be a bidirected graph. Let* {*uα, vβ* }*be a pair of vertex-sides with distinct u, v* ∈*V* (*G*) *and α, β* ∈ {+,−}. *Then {uα, vβ }is an* ultrabubble *if:*

a. separable: *the graph created by splitting uα and vβ contains a separate component X* ⊆*G containing u and v but not u*^′^ *and v*^′^. *We call X the* ultrabubble component *of {uα, vβ}*.
b. tipless: *no vertex in V* (*X*) \ {*u, v*} *is a tip*.
c. acyclic: *X is acyclic*.
d. minimal: *no vertex-side wγ with vertex w* ∈ *X* \ {*u, v*} *is such that* {*uα, wγ*} *and* 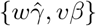*are separable*.

The *interior* of a separable pair of vertex-sides *{uα, vβ}* with component *K* is the vertex set *V* (*K*) \ {*u, v*}. A *trivial ultrabubble* is an ultrabubble whose component has vertex set {*u, v* }.

Consider an ultrabubble {*uα, vβ*}with component *X*. Notice that if 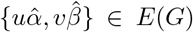 then this edge is clearly not contained in *X* due to the splitting operation. Thus, if we are to compute the ultrabubbles of a bidirected graph *G* by the computation of the superbubbles in an orientation of *G*, then we could miss those ultrabubbles with the “back-edge” (see Figure 2 and the Supplementary Material for an illustration). This is essentially the reason why we need weak superbubbles, and, indeed, we believe that the appropriate way to see ultrabubbles in directed graphs is through the notion of weak superbubbles (see Theorems 1 and 2).

### 2.2 Algorithm

To ensure the correctness of our algorithm we require the assumption that the input graph has at least one tip. We always make this assumption in what follows, and we present in the Supplementary Material the case where the graph has no tips, but at least one cutvertex.

Algorithm 1 performs a DFS starting from a tip *r* and maintains an array flipped[*v*] ∈{ `null, true, false`} for the vertices *v* ∈*V* (see Figure 3 (right) for what can go wrong if we do not start from a tip). This array serves the purpose of tracking which vertices are flipped and which vertices have already been visited. When arriving at a vertex-side *uα*, the DFS prioritizes exploring the edges of the form 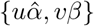, i.e., it first expands opposite-signed vertex-sides from the one at *u* when it was stacked. Moreover, when reaching an unvisited vertex *v* via an edge *uα, vβ, v* is flipped if and only if *α* = *β*, so flipped is updated as well as the edges of the graph incident to *v*. As a result, the traversed edge becomes directed (see Figure 3).

**Figure 3.**
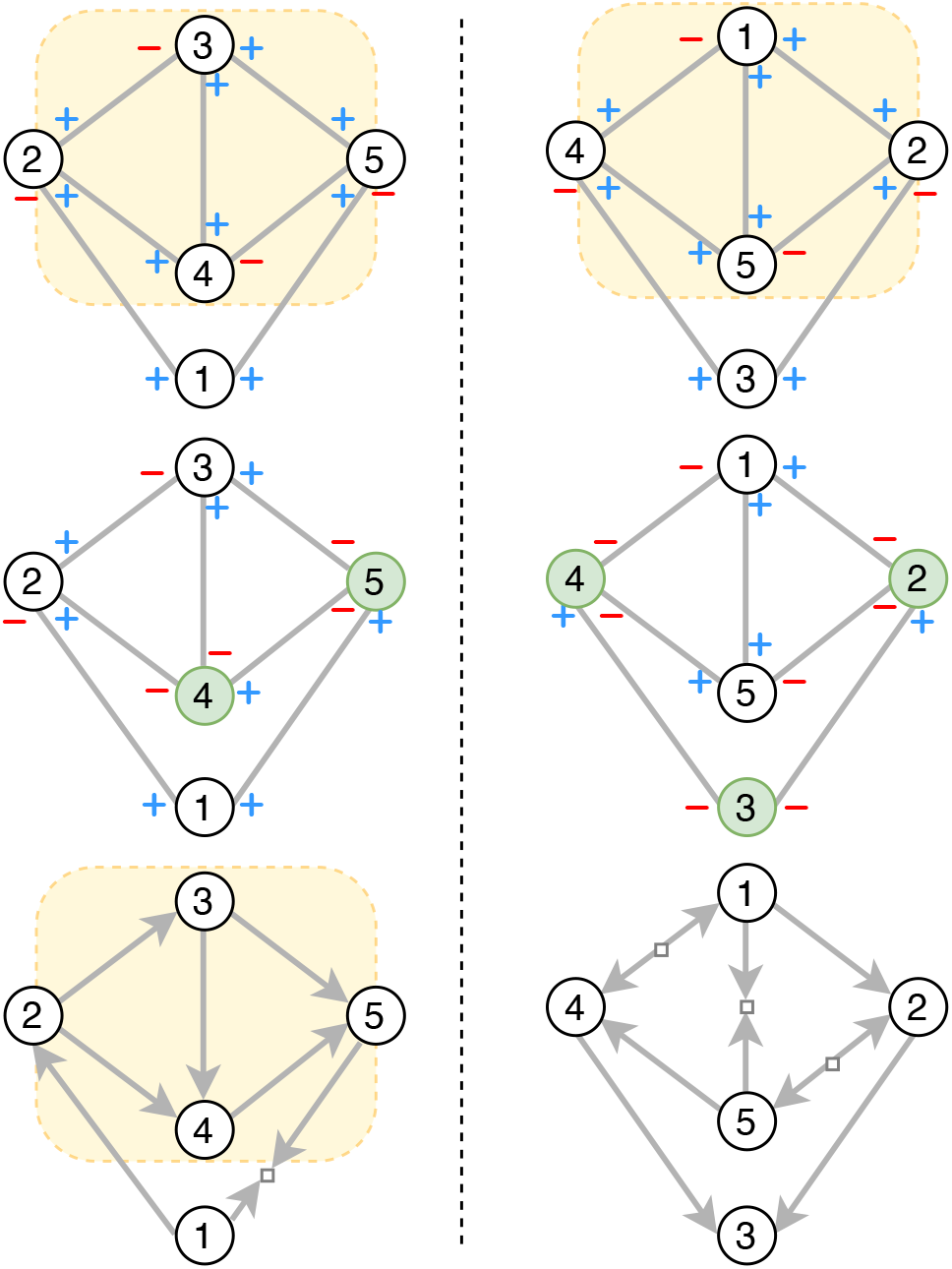
Example of the orientation algorithm on an ultrabubble. The left and right panels show the same graph, but with vertices numbered according to different DFS orders (both starting from vertex 1). On the left, vertex 1 is a tip, while on the right it is a non-tip vertex in the interior of an ultrabubble. **Left column**. Top: An ultrabubble 2+, 5+. Middle: The same ultrabubble after flipping the signs at vertices 4 and 5 (shown in green). The ultrabubble structure is preserved, and all its edges now have opposite signs. Bottom: The directed graph obtained by transforming each edge with opposite signs{*u*+, *v*−} into a directed edge from *u* to *v*. Under this transformation, the ultrabubble {2+, 5+} becomes the (weak) superbubble (2, 5). If the DFS had proceeded from vertex 1 immediately to the vertex now numbered 5, the resulting directed graph would instead contain the (weak) superbubble (5, 2), with all edges directed from right to left. **Right column**. Starting from vertex 1 and following the DFS order that determines the vertex numbering, vertices 2, 3, and 4 must be flipped. However, after these flips it is no longer possible to make all edges of the ultrabubble have opposite signs, which introduces sources and sinks in its interior. Consequently, the ultrabubble {4+, 2+}does not correspond to any superbubble in the resulting directed graph.

If later during the search the algorithm scans an equal-signed edge whose endpoints are visited (i.e., a “conflict edge”), this edge is subdivided with a fresh vertex *w* ∈ *Z* such that its two vertex-sides are both “−” if the conflict edge is plus-plus or they are both “+” if the conflict edge is minus-minus (see Figure 4).

**Figure 4.**
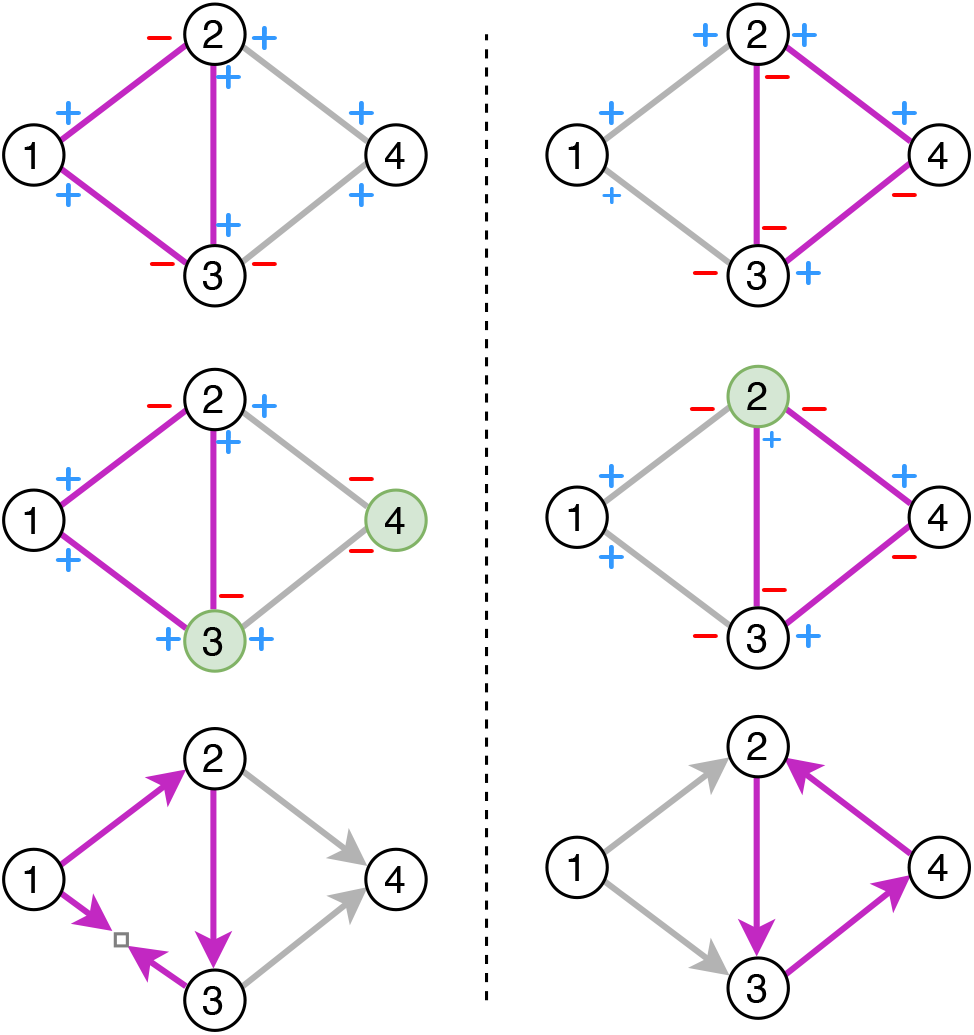
Example of the orientation algorithm on graphs that are not ultrabubbles. On the left column, we have a bidirected graph with a cycloid in violet, where only vertex 1 has the same sign at the edges incident to it. On the right column, we have a bidirected graph with a cycloid always alternating signs, also in violet. On the top row, we have the original graphs, with vertices numbered in a DFS order. On the middle row, we have the graph where the vertices in green have flipped signs. For the left-hand side graph, the same-sign edge {2+, 3+} requires flipping vertex 3, but then the original edge{ 1+, 3+}cannot be made to have distinct signs because 1 has already been processed and cannot be flipped anymore. As such, we add a fresh vertex in the middle of it, with edges oriented towards the tip. On the right-hand side graph, only vertex 2 needs to be flipped, and no vertices need to be introduced to obtain an equivalent bidirected graph where every edge has opposite signs. The cycle in the original graph maps to the directed cycle in violet.

We shall prove that no tip introduced during the DFS is contained in the vertex set of an ultrabubble component and that flipping vertices preserves the set of walks of the graph. These facts together will allow us to show the desired equivalence between ultrabubbles in the original instance and superbubbles in the graph produced by the reduction (with the sole exception that there can be new superbubbles ending or starting at tips introduced by the algorithm, but we do not report these as they do not correspond to ultrabubbles).

In what follows, we denote by *G* the input bidirected graph, by *D* the directed graph produced by Algorithm 1, and by *G*^⋆^ the bidirected graph that undergoes flips and addition of fresh vertices during Algorithm 1. At the beginning of this algorithm, the vertex set of *G*^⋆^ (implicitly) coincides with the vertex set of *G*, and the edge set of *G*^⋆^ (which is denoted by *E*^⋆^ in Algorithm 1) coincides with the edge set of *G*.

#### Algorithm 1

Orienting a bidirected graph

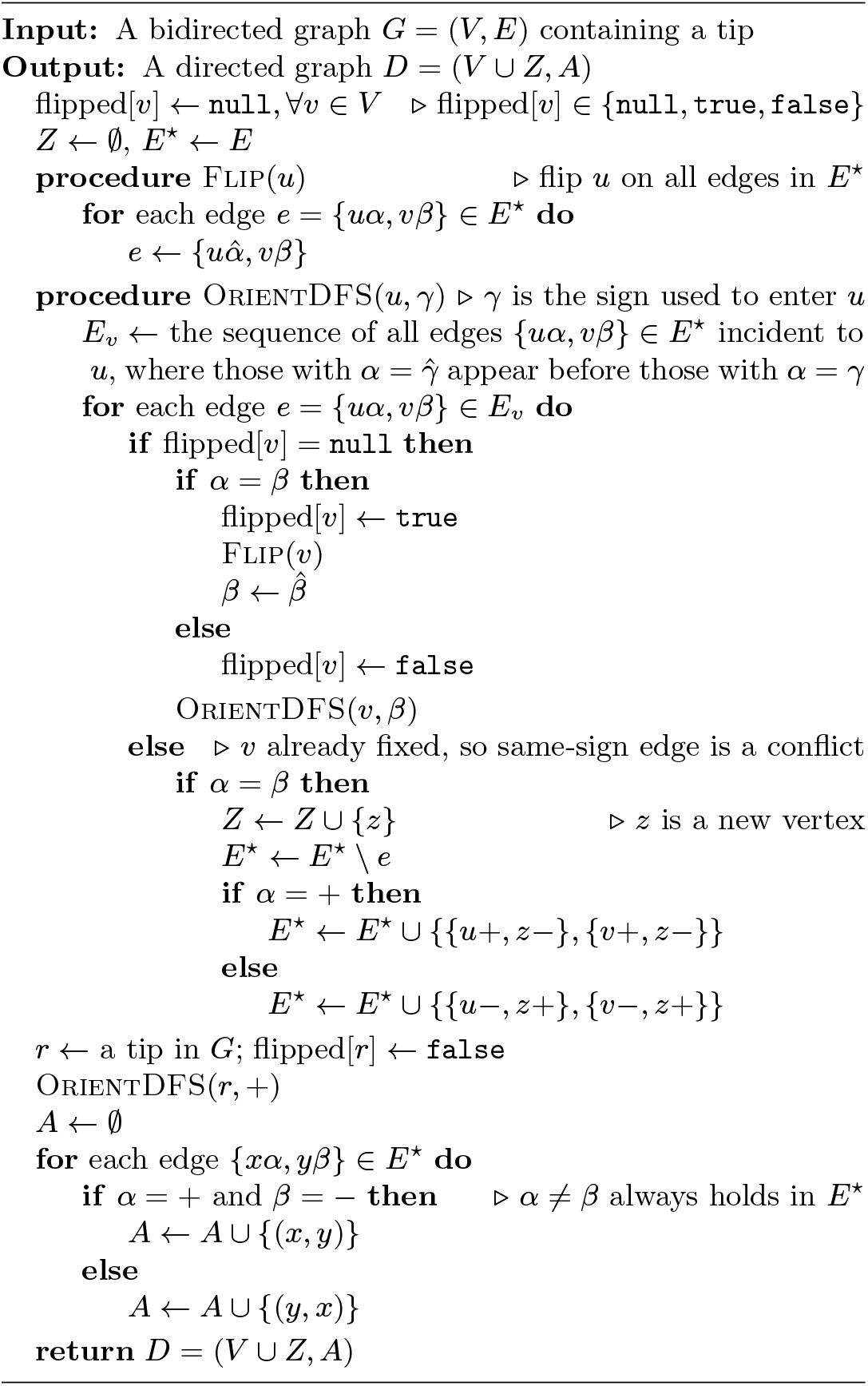

### 2.3 Correctness and complexity

Proofs of the following lemmas can be found in the Supplementary Material.

#### Lemma 1

*Let G be a bidirected graph, let u V* ∈ (*G*) *be a vertex, and let G*^′^ *be the graph obtained from G by flipping u. Let W be a sequence of edges in G and let W* ^′^ *be the same sequence of edges in G*^′^. *Then W is a walk in G if and only if W* ^′^ *is a walk in G*^′^.

The next lemma follows directly by examining Algorithm 1.

#### Lemma 2

*Algorithm 1 only changes G*^⋆^ *by flipping vertices or by adding tips via subdivision of edges*.

#### Lemma 3

*At the end of Algorithm 1 every edge of G*^⋆^ *has opposite signs in its vertex-sides, that is, every edge of G*^⋆^ *becomes directed*.

#### Lemma 4

*Let G be a bidirected graph and let* {*uα, vβ* } *be pair of vertex-sides with u≠v satisfying conditions (a), (b), and (c) of ultrabubbles (Definition 2). Let X denote the component of*{*uα, vβ*}. *Then uα, vβ is an ultrabubble if and only if no vertex in the interior of X is a u-v cutvertex in X*.

#### Corollary 1

*The underlying undirected graph of an ultrabubble component is biconnected*.

In what follows we assume implicitly that Algorithm 1 always produces a directed graph as output (e.g., in the assumptions of some statements). This raises no issues due to Lemma 3.

#### Lemma 5

*Let G be a bidirected graph and let {uα, vβ} be an ultrabubble with ultrabubble component X. Then Algorithm 1 does not subdivide any edge of X*.

#### Corollary 2

*Ultrabubble components are digraphic*.

#### Theorem 1

(Ultrabubble ⇒Weak superbubble). *Let G be a bidirected graph and let D be the directed graph produced by Algorithm 1. If {uα, vβ} is an ultrabubble of G then* (*u, v*) *or* (*v, u*) *is a weak superbubble of D*.

*Proof*. Let *X* be the component of the ultrabubble {*uα, vβ}*. Then no edge of *X* is subdivided with a tip by Lemma 5. Thus Lemma 2 implies that the only changes caused by Algorithm 1 to *X* were flipping vertices, and so *X*^⋆^ := *G*^⋆^[*V* (*X*)], *B* := *D*[*V* (*X*)], and *X* have the same vertex set. Moreover, Lemma 1 thus implies that *X* and *X*^⋆^ have the same set of walks. Recall that every edge of *G*^⋆^ has vertex-sides of opposite signs due to Lemma 3, and so *X*^⋆^ witnesses that *X* is digraphic.

To show reachability, first notice that a maximal bidirected path *p* in *X* starting at *u* witnesses that *X* has a *uα*-*vβ* bidirected path: otherwise, the last vertex *x v* on this path is entered, say, with sign *γ* ∈ {+, −}, and since *x* is not a tip any vertex in 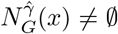 is already in *p* due to its maximality, but extending *p* with an edge from *x* to any such a vertex creates a cycloid, a contradiction. Thus *X*^⋆^ has a *uα*-*vβ* path and hence *B* has a directed path from *u* to *v* or from *v* to *u*. Assume without loss of generality in the remainder of the proof that *B* has a directed path from *u* to *v*. Notice that whenever 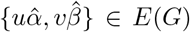then this edge is not subdivided in *G*^⋆^ and becomes the edge *vu* in *D*.

For acyclicity we have that *X* is acyclic and thus so is *X*^⋆^. Since *B*− *vu* is an orientation of *X*^⋆^, *B vu* is acyclic. This in turn implies that *u* is the source and *v* is the sink of *B*−*vu*.

We now argue on the matching condition. Every vertex *y* ∈ *V* (*X*) lies in some *u*-*v* path in *B* (consider, e.g., a longest path in *B*− *vu* containing *y*; the first and last vertices on this path are *u* and *v*, respectively, since *B*− *vu* is a directed acyclic graph with unique source *u* and unique sink *v*). If we show that every out-neighbor of *u* and every in-neighbor of *v* in *D* is a vertex of *B* we are done (i.e., so that the out-reachability of *u* avoiding *v* and the in-reachability of *v* avoiding *u* are confined to *B*). Since *X* is an ultrabubble component, by the separability prop-erty we have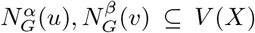. Since flipping vertices only swaps in-neighborhoods with out-neighborhoods, the same neighborhood containment holds in *G*^⋆^ up to flipping *α* and *β* so that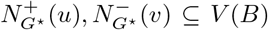. Therefore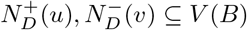.

To see minimality suppose for a contradiction that *B* has a vertex *w* witnessing its non-minimality. If *vu ∉E*(*B*) then *B* is a superbubbloid that is not a superbubble and so *w* is a *u*-*v* cutvertex in *B* (to see why, see (Sena et al., 2025); essentially, an analogous statement to that of Lemma 4 holds in the context of superbubbles). So *w* is a *u*-*v* cutvertex in *X*^⋆^ and thus of *X* too, a contradiction to Lemma 4. Otherwise *vu ∈ E*(*B*). Since (*u, w*) is a weak superbubbloid, say with graph *B*^′^, then *vu ∉ E*(*B*^′^) because *v ∉V* (*B*^′^). Moreover, *wu ∉ E*(*B*^′^) for otherwise *wu ∉E*(*B*) and thus *B* has a cycle without *vu*, a contradiction to the definition of weak superbubbloid containing the edge *vu*. So (*u, w*) is a super-bubbloid and by a symmetrical argument we can conclude that (*w, t*) is a superbubbloid. Thus *B – vu* is a superbubbloid with *w* witnessing its non-minimality and hence *w* is a *u*-*v* cutvertex in *B*−*vu* (see (Sena et al., 2025)). Since *X*^⋆^ does not contain the edge {*v*+, *u*−} due to the splitting operation, *w* is a *u*-*v* cutvertex in *X*^⋆^ and so *w* is a *u*-*v* cutvertex in *X*, a contradiction to Lemma 4.

Thus, (*u, v*) is a weak superbubble in *D* if *u* is the source of *B* or (*v, u*) is a weak superbubble in *D* if *v* is the source of *B*.

#### Theorem 2

(Weak superbubble ⇒Ultrabubble). *Let G* = (*V, E*) *be a bidirected graph and let D be the directed graph produced by Algorithm 1. Let* (*u, v*) *be a weak superbubble of D with u, v* ∈ *V*. *Let α, β* ∈ {+, −} *be such that if* flipped[*u*] = *`false` then α* = +, *otherwise α* = −, *and if* flipped[*v*] = *`false` then β* = −, *otherwise β* = +. *Then* {*uα, vβ*} *is an ultrabubble of G*.

*Proof*. Let *B* be the component of the weak superbubble (*u, v*). Then *V* (*B*) \{*u, v }*contains no sources or sinks of *B* by the matching condition and therefore Algorithm 1 did not introduce tips in the edges of *G*[*V* (*B*)] via subdivisions. Thus, by Lemma 2, the DFS only altered *G*[*V* (*B*)] by flipping some of its vertices and so *G*^⋆^[*V* (*B*)] and *G*[*V* (*B*)] have the same reachability relation by virtue of Lemma 1. Notice also that *X* := *G*[*V* (*B*)], *X*^⋆^ := *G*^⋆^[*V* (*B*)] and *B* have the same vertex set.

We argue on the separability. Notice that any directed path in *D* from a vertex not in *B* to a vertex in *B* contains *u* since *u* is the entry of the weak superbubble, and similarly, any path from a vertex in *B* to a vertex not in *B* contains *v* since *v* is the exit of the weak superbubble (see (Gärtner et al., 2018; Gärtner and Stadler, 2019) for a definition of weak superbubble exposing these properties). Since *B* has a *u*-*v* directed path and trivially 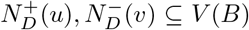 and 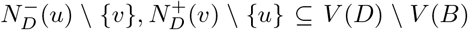, it follows that {*u*+, *v*− }is separable in *G*^⋆^ with component *X*^⋆^. Since *X*^⋆^ was obtained from *X* only by flipping vertices, it follows that {*uα, vβ*}is separable in *G*.

We now argue on the acyclicity. Since *B*− *vu* is acyclic so is *X*^⋆^ (notice that *X*^⋆^ does not contain the potential bidirected edge {*v*+, *u*−} due to the splitting operation). Thus *X* is acyclic because *X* does not contain the potential bidirected edge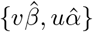.

We are left to show minimality. Notice that *X* ⊆ *G* is the component containing *u* and *v* and not *u*^′^ and *v*^′^ after splitting *uα* and *vβ* in *G*, and that *X*^⋆^ ⊆ *G*^⋆^ is the component containing *u* and *v* and not *u*^′^ and *v*^′^ after splitting *u*+ and *v*− in *G*^⋆^. Suppose for a contradiction that {*uα, vβ* }is not minimal, so *X* has a a *u*-*v* cutvertex *w* in *X* by Lemma 4. Thus *w* is a *u*-*v* cutvertex in *X*^⋆^ since subdividing edges and flipping vertices does not create an undirected *u*-*v* path in *X*^⋆^ avoiding *w*. So if *vu ∉ E*(*B*) then *B* is a superbubble and *w* is a *u*-*v* cutvertex in *B* since *B* has the same set of undirected paths as *X*^⋆^, a contradiction to the minimality of (*u, v*) (see Sena et al. (2025)). Otherwise, if *vu* ∈*E*(*B*) then by the same argument used to show minimality in the proof of Theorem 1 we can conclude that *w* is a *u*-*v* cutvertex of *B*− *vu* and thus (*u, w*) is a superbubbloid, a contradiction to the minimality of (*u, v*).

We can conclude that {*uα, vβ*} is an ultrabubble of *G*.

#### Theorem 3

*We can compute all ultrabubbles of a bidirected graph G with at least one tip, or a cutvertex, in time O*(|*V* (*G*)| + |*E*(*G*)|).

## 3 Implementation and Experiments

### 3.1 Implementation

We implemented the new algorithm in our `BubbleFinder` tool (Sena et al., 2025), available at www.github.com/algbio/BubbleFinder, as a new `ultrabubbles` subcommand. We used the C++ implementation of the weak superbubble algorithm by Gärtner and Stadler (2019) from www.github.com/Fabianexe/clsd. Tips are identified during parsing and serve as starting points for the orientation DFS; for tipless components, cutvertices are identified via Tarjan’s algorithm (Tarjan, 1972). The oriented graph is passed to the CLSD weak superbubble algorithm of Gärtner and Stadler (2019). In practice, few conflict vertices are introduced (Supplementary Material, Table 2): 2,691 on HPRC v1.1 (92.9M nodes) and 244,265 on v2.0 (148.3M nodes); the `vg` graphs, being already directed, produce none.

**Table 1.** Performance comparison of `vg` snarls and `BubbleFinder`, on HPRC graphs in GBZ format. Times are wall-clock (single-threaded), formatted as h:mm:ss or m:ss. Memory is peak RSS in GiB. Both tools use the same GBZ parsing library (`gbwtgraph`). Speedup is computed excluding shared GBZ parsing time.

| Type | Dataset | vg snarls |  | BubbleFinder |  | Parsing | Speedup |
| --- | --- | --- | --- | --- | --- | --- | --- |
|  |  | Time | Mem (GiB) | Time | Mem (GiB) | (GBZ, s) | excl. parsing |
| MC | HPRC v1.1 Chr. X | 2:02 | 5.1 | 0:15 | 1.7 | 0:10 | 24.74× |
|  | HPRC v1.1 Chr. Y | 0:32 | 1.7 | 0:05 | 0.56 | 0:04 | 18.97× |
|  | HPRC v1.1 CHM13 (47 indiv.) | 37:23 | 66.3 | 3:54 | 14.3 | 2:34 | 25.99× |
|  | HPRC v2.0 CHM13 (232 indiv.) | 1:12:13 | 101.8 | 9:40 | 24.8 | 7:04 | 25.09× |

**Table 2.**
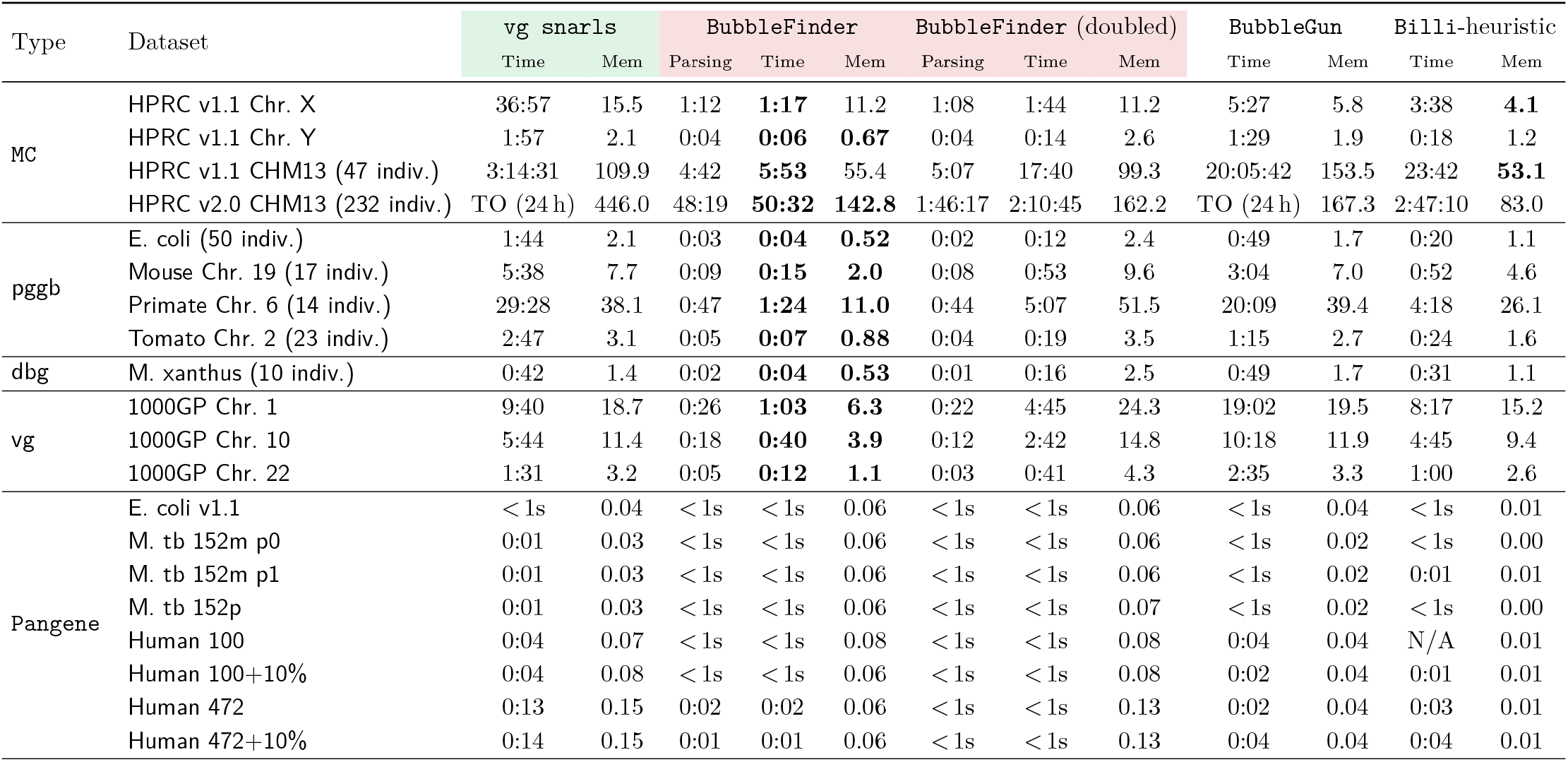
Performance comparison on GFA graphs across five tools computing bubble-like structures. Times are wall-clock (single-threaded), formatted as h:mm:ss or m:ss. Memory is peak RSS in GiB. For `BubbleFinder` and `BubbleFinder` (doubled), the Parsing column reports the parsing part of the runtime (including on-the-fly construction of the internal graph representation) and is included in the Time column. For the doubled mode, this involves building a graph with twice as many vertices. `BubbleFinder` (doubled) uses the doubled-graph method for ultrabubble detection. Best time and memory per dataset are in **bold** (excluding parsing-only columns and datasets where all tools take under one second). TO = exceeded time limit; N/A = tool does not run because the graph is tipless.

`BubbleFinder` supports GBZ and GFA input formats; for GBZ, it uses the same `gbwtgraph` library (Sirén and Paten, 2022) as `vg. BubbleFinder` also supports the `–T` flag of `vg` to additionally output trivial, i.e. single-edge, ultrabubbles. `BubbleFinder` also supports finding and merging (weak) superbubbles from the doubled directed graph constructed from the bidirected graph, which is (conceptually) implemented by `BubbleGun` (see also the Supplementary Material).

As an additional feature in `BubbleFinder`, we also implement the computation of the tree hierarchy of ultrabubbles. This is based on the superbubble decomposition computed by Gärtner and Stadler (2019), which we transform into an ultrabubble decomposition by removing superbubbles whose entrance or exit corresponds to a newly introduced tip and connecting its children to their parent in the tree hierarchy. The resulting *ultrabubble decomposition* forms a rooted forest, see the two nested ultrabubbles in Figure 5 (akin to the snarl decomposition of Paten et al. (2018), only that snarls can also contain cycles or tips).

**Figure 5.**
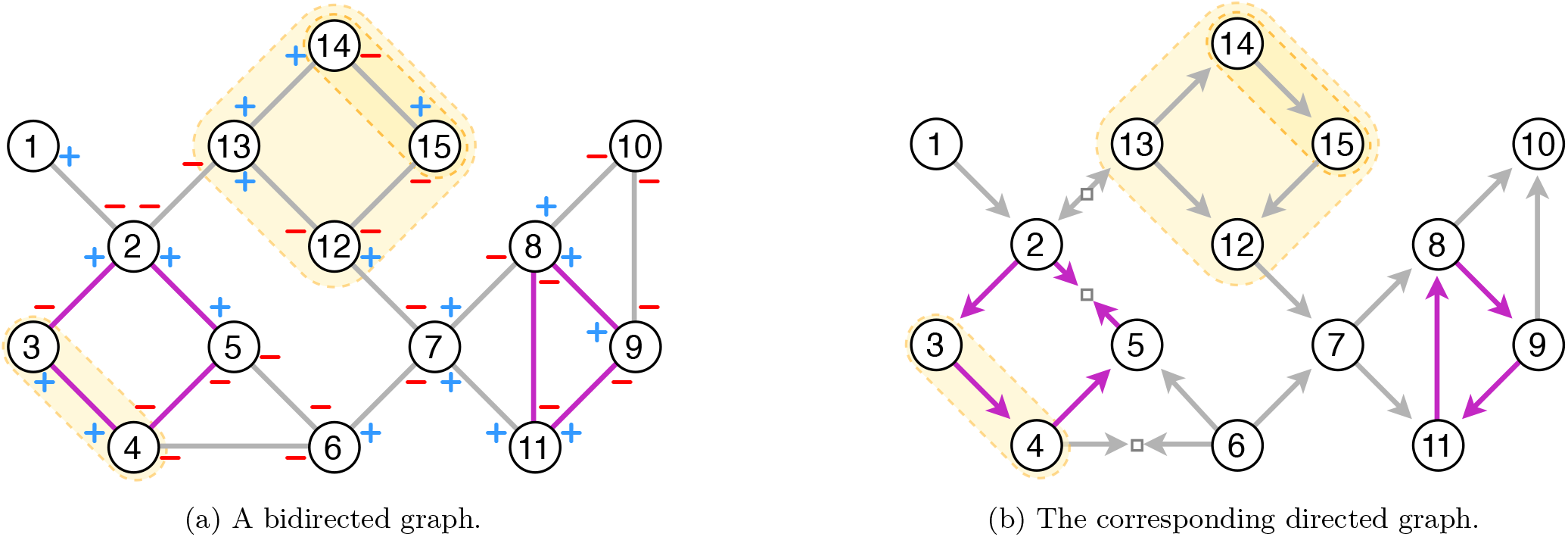
A bidirected graph and the directed graph constructed by Algorithm 1. Highlighted in yellow, the graphs contain three ultrabubbles ({3+, 4+, 13+, 12, 14, 15+ }) and superbubbles ((3, 4), (13, 12), (14, 15)), respectively (no proper weak superbubbles are shown in this figure). Note that ultrabubble {13+, 12−} includes ultra-bubble {14−, 15+},and superbubble (13, 12) includes superbubble (14, 15). Highlighted in purple, the bidirected graph contains a cycloid {2+, 3−}, {3+, 4+}, *{*4−, 5−}, *{*5+, 2+} with exception at vertex 2, which introduces a sink in the directed graph between vertices 2 and 5, and a cycloid {8+, 9+}, *{*9−, 11+}, {11−, 8−} without exceptional vertex, which becomes a directed cycle.

### 3.2 Experimental results

#### Datasets

We perform experiments on five families of pangenome graphs: (i) `MC`: Human Pangenome Reference Consortium (HPRC) Minigraph-Cactus `(MC)` graphs, downloaded from the HPRC data portal, ^4^ including HPRC v1.1 (47 individuals) and v2.0 (232 individuals); (ii) `pggb`: E. coli, Primate Chr. 6, Tomato Chr. 2 and Mouse Chr. 19 are pangenome graphs constructed with `pggb` (Garrison et al., 2024); (iiii) `dbg`: a de Bruijn graph constructed from 10 individuals of M. xanthus, used in the evaluation of `BubbleGun` (Dabbaghie et al., 2022); (iv) `vg`: Chromosome 1 |10| 22 are variant graphs for human chromosome 1 |10| 22 constructed from the (∼ 6.5 millions) variant calls from phase 3 of 1000 Genomes Project Consortium (2015). Accession codes and URLs for the graphs in the `pggb` and `vg` datasets are in (Sena et al., 2025). (v) Pangene: complex gene graphs by Li et al. (2024), including E. coli, M. tb and Human pangene graphs, used in the evaluation of `Billi` (Bhat et al., 2025). See the Supplementary Material (Table 2) for statistics on the size of all these graphs, and the numbers of tips and cutvertices.

#### Tools

We compare `BubbleFinder` with the ultra-bubble finding algorithm by Paten et al. (2018), available in `vg` via the subcommand `snarls`. We also compare with `BubbleGun` (Dabbaghie et al., 2022), and with the panbubble heuristic of `Billi` (Bhat et al., 2025) (we did not use the exact algorithm implemented in `Billi`, because as observed in (Bhat et al., 2025), the exact one does not scale to large pangenome graphs, and it has the same behavior as the heuristic version on the smaller graphs tested in (Bhat et al., 2025)). We used `vg` v1.72.0, `BubbleGun` v1.2.0, and `Billi` v1.0.^5^ Our snake-make pipeline for the experiments is available at github. com/algbio/BubbleFinder-experiments/. All our experiments ran on Intel(R) Xeon(R) CPU E7-8890 v4 @ 2.20GHz machines with 1024 GB of RAM.

## Results

A detailed comparison of ultrabubble counts across all tools is given in the Supplementary Material (Table 1). On all tested datasets, `BubbleFinder` reports the same number of ultrabubbles as `vg` (both trivial and non-trivial). Moreover, also the mode of `BubbleFinder` of computing and merging superbubbles in the doubled graph reports the same number of bubbles in all the tested graphs. However, this generally uses more than double the time and memory of the orientation-based mode, and to the best of our knowledge there exists no proof that this approach correctly computes the set of ultrabubbles.

On the pangenome graphs `MC, pggb` and `vg`, the difference between snarls (including trivial) and ultrabubbles (including trivial) is very small. Likewise, the difference between the number of panbubbles computed by `Billi`-heuristic and non-trivial ultrabubbles is also very small. While the non-ultrabubble subgraphs may capture relevant structures, on these graphs ultrabubbles appear to cover a very large proportion of snarls and panbubbles. On the `dbg` graph, there are only three more panbubbles than non-trivial ultrabubbles and on the `vg` graphs all numbers match. On the `Pangene` graphs the bubble structures are more different, for example there are even more panbubbles than snarls on some datasets, and as (Bhat et al., 2025) already showed, panbubbles do capture relevant biological events in such cases. However, these graphs are much smaller than pangenome graphs (having at most 42,000 edges). Moreover, notice this dataset contains one tipless graph where Billi cannot run (since it assumes graphs with at least one tip), while this can still be handled by `BubbleFinder` because it has a cutvertex.

In Table 1 we show the comparison between `vg` and `BubbleFinder` on the HPRC graphs in GBZ compressed format (Sirén and Paten, 2022) with the -T flag. This format is designed to be parsed much more efficiently than GFA, as we also observe in the “Parsing” column of Table 1 (see also Table 2 for GFA runtimes). Excluding the shared GBZ parsing time, `BubbleFinder` achieves speedups of 19–26× over `vg` on the HPRC graphs. On HPRC v2.0, after parsing, `BubbleFinder` completes in under 3 minutes while `vg` requires more than one hour and four times more RAM (101.8 GiB vs 24.8 GiB). In fact, also on the smaller datasets, the memory consumption of `BubbleFinder` is on par with or lower than that of `vg`. The 25× speedup combined with the 4-fold reduction in memory footprint highlights the practical advantages of our linear-time orientation approach for population-scale pangenomics. Moreover, we tried to convert the other datasets in GBZ format, but the haplotypes in the GFA file did not cover all edges present in it, and thus we could not obtain equivalent GBZ graphs, and hence omitted the results.

Since `BubbleGun` and `Billi` do not support GBZ input, in Table 2 we compare all tools on graphs in GFA format. For completeness, we also compare with `vg` on these graphs with the -T flag, same as for `BubbleFinder`. On the v1.1 HPRC graph, our method is 200× faster than `BubbleGun`, and on the HPRC v2.0, `BubbleFinder` completed in under 51 minutes, while `BubbleGun` timed out. Moreover, `BubbleGun` seems to report a few cases of bubbles that are not ultrabubbles, see the Supplementary Material. `BubbleFinder` is faster also than `Billi`-heuristic (4 ×on HPRC v1.1 and 3×on HPRC v2.0), but note that `Billi` computes more complex structures.

## 4 Conclusion

While bidirected graphs are more expressive than directed graphs—since they naturally capture the reverse complementarity of DNA—we showed that, for the purpose of computing ultrabubbles, they can be oriented to yield an ordinary directed graph (without the need to double the graph, and with a very small number of additional vertices, in practice). Under this orientation, the more general problem of finding ultrabubbles reduces to computing weak superbubbles. This leads to substantial speedups compared to computing (ultra)bubbles with other tools.

Our orientation algorithm raises the broader question of which other variation structures on bidirected graphs can be reduced to directed graphs. Natural candidates include bibubbles (Li et al., 2024) and panbubbles (Bhat et al., 2025), for which no linear-time algorithms are currently known. Any such orientation reductions would likely need to be substantially more sophisticated than the one presented here, reflecting the greater structural complexity of these objects (e.g., they can contain cycles, but require a stronger reachability property on the bubble vertices). Moreover, unlike for ultrabubbles, there are no known directed counterparts for bibubbles or panbubbles.

## Acknowledgments

We thank Massimo Cairo and Romeo Rizzi for very fruitful discussions. This work was co-funded by the European Union (ERC, SCALEBIO, 101169716). Views and opinions expressed are however those of the author(s) only and do not necessarily reflect those of the European Union or the European Research Council. Neither the European Union nor the granting authority can be held responsible for them. Juha Harviainen was supported by the Research Council of Finland, Grant 351156 and by the Helsinki Institute of Information Technology (HIIT).

## Footnotes

1 https://humanpangenome.org/hprc-data-release-2/

2 https://humanpangenome.org/release-timeline/

3 A cycloid is a closed walk, and a closed walk that is not a cycloid can be short-circuited to obtain a cycloid.

4 https://data.humanpangenome.org/alignments/

5 https://github.com/at-cg/billi, commit a77892e.

## References

1000 Genomes Project Consortium. A global reference for human genetic variation. Nature, 526(7571):68, 2015.

T. Abrishami, N. Bowler, A. Joó, F. Reich, and Q. Tao. A structure theorem for rooted connectivity in bidirected graphs. arXiv preprint arXiv:2509.23394, 2025.

S. G. Bhat, D. Mahajan, and C. Jain. Billi: Provably accurate and scalable bubble detection in pangenome graphs. bioRxiv, 2025. URL 10.1101/2025.11.21.689636.

F. Dabbaghie, J. Ebler, and T. Marschall. BubbleGun: enumerating bubbles and superbubbles in genome graphs. Bioinformatics, 38(17):4217–4219, 07 2022.

D. Fasulo, A. Halpern, I. Dew, and C. Mobarry. Efficiently detecting polymorphisms during the fragment assembly process. In ISMB, pages 294–302, 2002.

E. Garrison et al. Variation graph toolkit improves read mapping by representing genetic variation in the reference. Nature biotechnology, 36(9):875–879, 2018.

E. Garrison et al. Building pangenome graphs. Nature Methods, 21(11):2008–2012, Nov 2024.

F. Gärtner and P. F. Stadler. Direct superbubble detection. Algorithms, 12(4):81, 2019.

F. Gärtner, L. Müller, and P. F. Stadler. Superbubbles revisited. Algorithms for Molecular Biology, 13(1):16, 2018.

N. Kita. Bidirected Graphs I: Signed General Kotzig-Lovász Decomposition. arXiv preprint arXiv:1709.07414, 2017.

M. Kolmogorov et al. metaFlye: scalable long-read metagenome assembly using repeat graphs. Nat. Methods, 17(11):1103–1110, 2020.

H. Li, M. Marin, and M. R. Farhat. Exploring gene content with pangene graphs. Bioinformatics, 40(7):btae456, 2024.

W.-W. Liao et al. A draft human pangenome reference. Nature, 617(7960):312–324, 2023.

I. Minkin and P. Medvedev. Scalable multiple whole-genome alignment and locally collinear block construction with SibeliaZ. Nature communications, 11(1):6327, 2020.

T. Onodera, K. Sadakane, and T. Shibuya. Detecting superbubbles in assembly graphs. In WABI 2013, pages 338–348. Springer, 2013.

B. Paten, J. M. Eizenga, Y. M. Rosen, A. M. Novak, E. Garrison, and G. Hickey. Superbubbles, ultrabubbles, and cacti. Journal of Computational Biology, 25(7):649–663, 2018.

A. Rahman and P. Medvedev. Assembler artifacts include misassembly because of unsafe unitigs and underassembly because of bidirected graphs. Genome Res., 32(9):1746–1753, 2022.

M. Rautiainen et al. Telomere-to-telomere assembly of diploid chromosomes with Verkko. Nat. Biotechnol., 41 (10):1474–1482, 2023.

F. Sena, A. Politov, C. Moumard, M. Cáceres, S. Schmidt, J. Harviainen, and A. I. Tomescu. Identifying all snarls and superbubbles in linear-time, via a unified SPQR-tree framework, 2025. URL https://arxiv.org/abs/2511.21919.

J. Sirén and B. Paten. GBZ file format for pangenome graphs. Bioinformatics, 38(22):5012–5018, 2022.

R. Tarjan. Depth-first search and linear graph algorithms. SIAM Journal on Computing, 1(2):146–160, 1972.

D. R. Zerbino and E. Birney. Velvet: algorithms for de novo short read assembly using de Bruijn graphs. Genome research, 18(5):821–829, 2008.

A. E. Zisis and P. Sætrom. Ultrabubble enumeration via a lowest common ancestor approach, 2026. URL https://arxiv.org/abs/2603.03909.

