## Supplementary Material for "Scalable computation of ultrabubbles in pangenomes by orienting bidirected graphs"

### Scalable computation of ultrabubbles in pangenomes by orienting bidirected graphs
### — Supplementary Material —

#### Contents

**1 Ultrabubbles in biedged graphs** **2**

**2 Ultrabubbles vs. snarls, panbubbles, bibubbles** **2**

**3 Computing (weak) superbubbles in the doubled directed graph** **4**

**4 Additional methods** **5**

4.1 Missing proofs . . . . . 5

4.2 Handling graphs with cutvertices . . . . . 9

**5 Graph statistics and bubble counts** **10**

### 1 Ultrabubbles in biedged graphs

Ultrabubbles were defined by Paten et al. (2018) in terms of biedged graphs. In Figure 1 we illustrate the equivalence between ultrabubbles in bidirected graphs and ultrabubbles in biedged graphs.

In Figure 1a we have the example from Figure 1 in the Main Paper, with an ultrabubble $\{3+, 6+\}$ in a bidirected graph $G$, shown as a yellow rounded rectangle. In Figure 1b we have the biedged graph equivalent to $G$. This is an undirected graph whose edges are either *black* or *gray*. This is obtained by adding a black edge labeled $v$ for every vertex $v$ of $G$, and adding gray edges between endpoints of black edges with the following convention: every "left" vertex of a black edge $v$ gathers the bidirected edges incident to $v-$, and every "right" vertex of a black edge $v$ gathers the bidirected edges incident to $v+$. In Figure 1b, we draw in yellow the vertices contained in the ultrabubble $\{3+, 6+\}$.

In Figure 1c we have an equivalent, but simpler, drawing of the biedged graph from Figure 1b, obtained by "flipping" the drawing of the black edges labeled 1, 5, 6, 7, 8. After flipping, the ultrabubble $\{3+, 6+\}$ can be visually identified more easily. Note that flipping these black edges is analogous to our flipping operation on bidirected graphs (flipping the signs of the bidirected edges incident to a vertex) that we used in Algorithm 1 in the Main Paper.

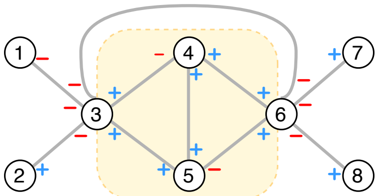


(a) A bidirected graph $G$

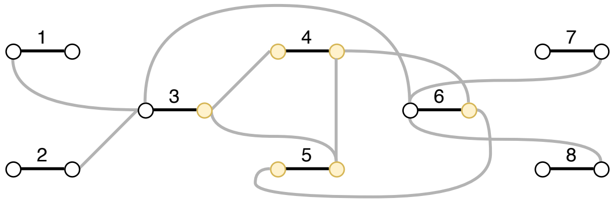


(b) A biedged graph obtained from $G$

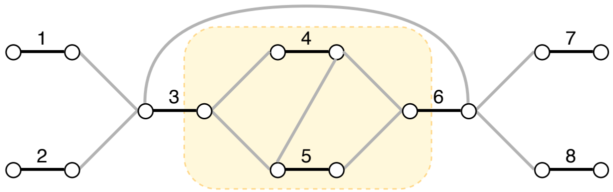


(c) An equivalent drawing of the biedged graph from Figure 1b

Figure 1: Example of an ultrabubble $\{3+, 6+\}$ in a bidirected graph and in an equivalent biedged graph.

### 2 Ultrabubbles vs. snarls, panbubbles, bibubbles

In this section, we give examples illustrating the differences between snarls (Paten et al., 2018), panbubbles (Bhat et al., 2025) / bibubbles (Li et al., 2024) and ultrabubbles (Paten et al., 2018); please refer to Figure 2. In this figure, on the left column, we have a snarl $\{3+, 6-\}$ which contains a cycle $\{9, 10, 11\}$ (in violet). Moreover, the vertices of this cycle do not reach $6-$, and thus $\{3+, 6-\}$ is also not a panbubble/bibubble.

On the middle column, we have the same graph as on the left column, except that there is an additional edge between vertices 9 and 5. Now, $\{3+, 6-\}$ is a panbubble / bibubble, since the presence of this edge make all vertices in the bubble reachable from $3+$ and reaching $6-$ (a walk from $3+$ to $6-$ containing all vertices of the bubble is shown in violet). The bubble $\{3+, 6-\}$ is also not an ultrabubble, because it still contains the cycle $\{9, 10, 11\}$.

On the right column, we have a similar graph as on the left column, only that the orientation of some edges are different. We have that $\{3+, 6-\}$ is an ultrabubble in this graph, since the bubble contains no cycle.

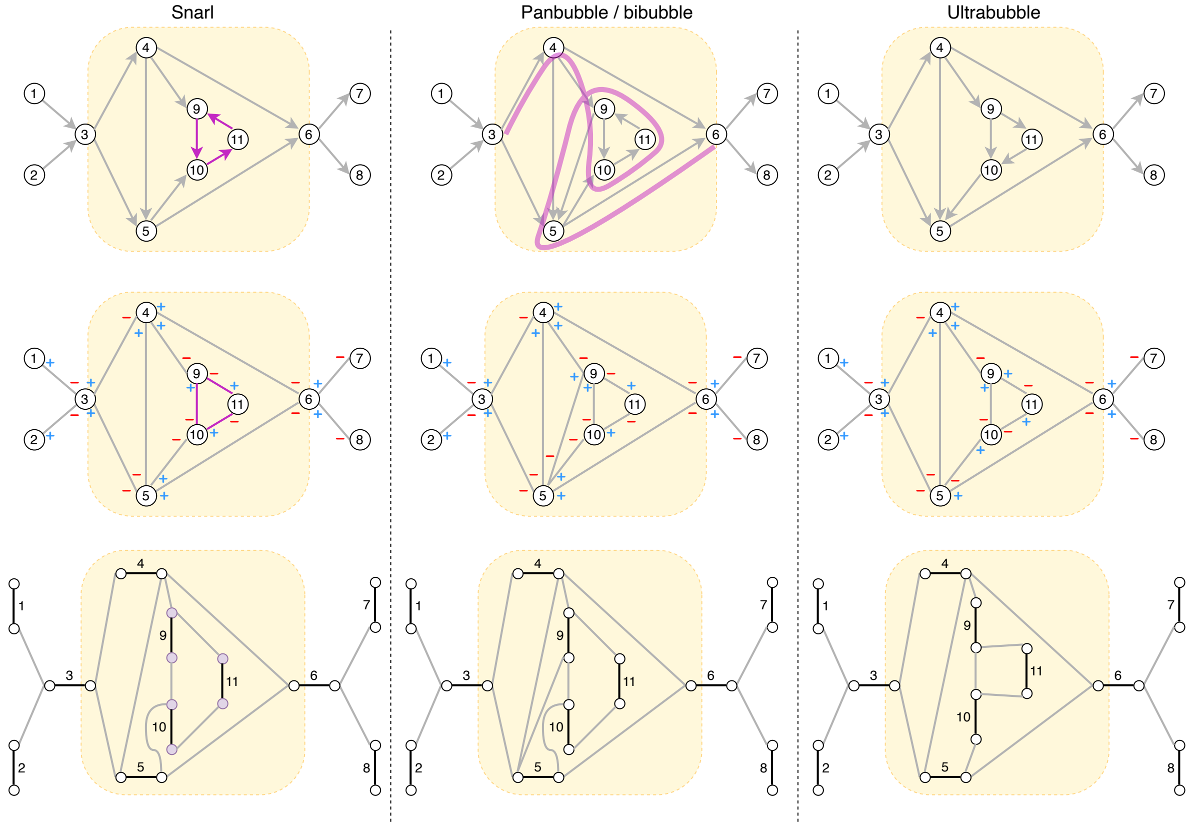


Figure 2: Examples of three graphs (one per column). Each graph is obtained starting from a directed graph (top row), which on the second row is illustrated as a bidirected graph with the convention that directed edges $(u, v)$ get converted to bidirected edges $\{u+, v-\}$. Each graph is also illustrated on the bottom row as a biedged graph, using the convention from Figure 1. Each column shows as a yellow rounded rectangle a snarl, a panbubble / bibubble and an ultrabubble, respectively.

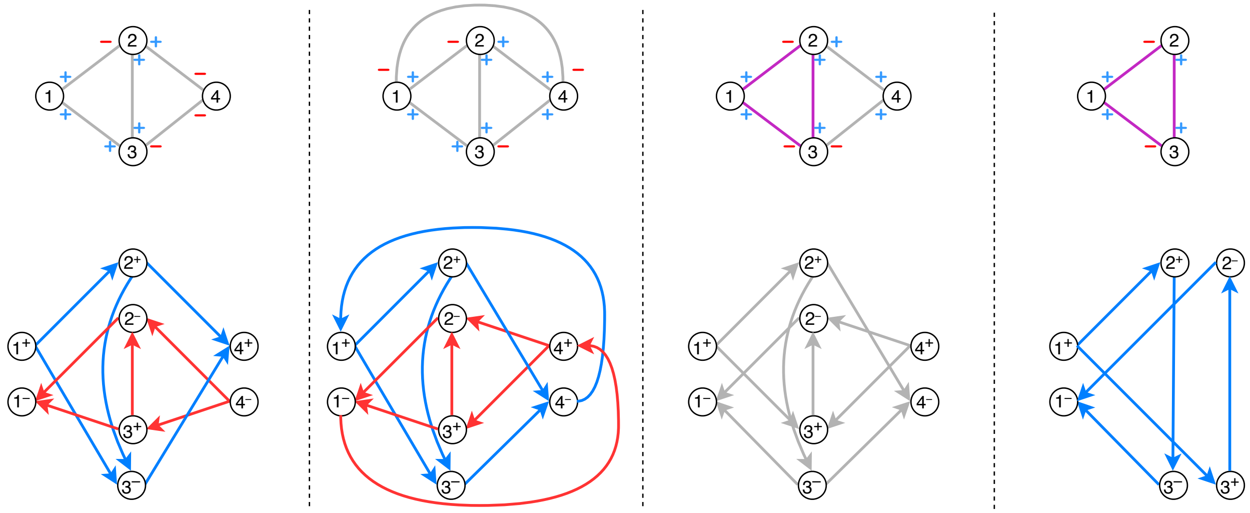

Figure 3: Example of the doubled graph applied to four bidirected graphs. The top row shows the bidirected graph, and the bottom row shows the doubled graph. In the graph on the first column (from the left), $\{1+, 4-\}$ is an ultrabubble, which then corresponds to superbubbles $(1+, 4+)$ and $(4-, 1-)$ (in blue, and red, respectively) in the doubled graph. On the second column, $\{1+, 4+\}$ is an ultrabubble (with the bidirected edge $\{1-, 4+\}$ outside the ultrabubble), which then corresponds to proper *weak* superbubbles $(1+, 4-)$ and $(4+, 1-)$ (in blue, and red, respectively) in the doubled graph. On the third column, the bidirected graph contains a cycle (in violet), and thus in the doubled graph $(1+, 4-)$ and $(4+, 1-)$ are no longer superbubbles. In the graph on the right-most column, the cycle in violet starting and ending at 1 leads to a (unique) superbubble $(1+, 1-)$ in the doubled graph, having the same vertex ID 1 of the bidirected graph.

#### 3 Computing (weak) superbubbles in the doubled directed graph

An approach to compute acyclic bubbles in bidirected graphs is to "double" the graph, which results in a directed graph, and compute (weak) superbubbles in this doubled directed graph. Conceptually, this approach is employed by the tool BubbleGun (Dabbaghie et al., 2022). More specifically, given a bidirected graph $G$, one constructs a directed graph $D$ having vertices $v+$ and $v-$ for every vertex $v$ of $G$. For every bidirected edge $\{u\alpha, v\beta\}$, we add directed edges in $D$ from vertex $v\alpha$ to $u\hat{\beta}$ and from $u\beta$ to $v\hat{\alpha}$. One can interpret each $v+$ and $v-$ as a *state*: $v+$ is "need to exit with $+$", and $v-$ is "need to exit with $-$". For example, if we have a bidirected edge $\{v+, u+\}$, the directed edge is from vertex $v+$ to vertex $u-$, because all walks must continue from vertex $u$ in the bidirected graph with an edge having sign $-$. Intuitively, vertex $u-$ (which is the state "need to exit with $-$") gathers these outgoing directed edges. Analogously, we also have the directed edge from $u+$ to $v-$.

The transformation of (weak) superbubbles in the doubled directed graph to "bubbles" in the bidirected graph is the following: every pair of symmetric (weak) superbubbles of the form $(v\alpha, u\beta)$ and $(u\hat{\beta}, v\hat{\alpha})$ (with $u \neq v$) are "merged" a bubble $\{v\alpha, u\hat{\beta}\}$ in the original bidirected graph. (Weak) superbubbles with $u = v$ are not reported. See Figure 3 for examples of bidirected graphs and their doubled directed graphs, and of this merging operation of superbubbles.

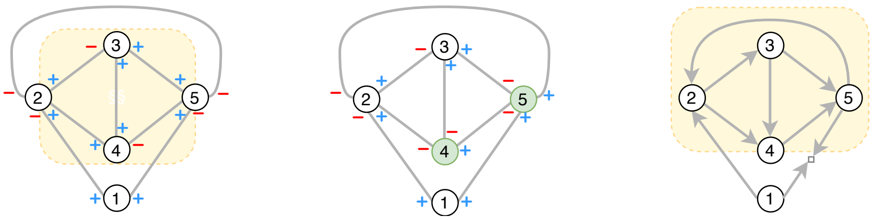

Figure 4: **An ultrabubble with an "opposite" edge between its endpoints translates to a weak superbubble.** On the left, we have an ultrabubble $\{2+, 5+\}$ such that the graph also has an edge $\{2-, 5-\}$ outside of the ultrabubble. The orientation algorithm performs flips as in the middle figure, and after orientation (in the right figure) the bidirected edge $\{2-, 5-\}$ becomes a directed edge $(5, 2)$. However, $(2, 5)$ is a weak superbubble, and not a superbubble.

### 4 Additional methods

In Figure 4 we show an example of an ultrabubble with an edge between its endpoints (which is outside of the ultrabubble), which in the directed graph becomes a weak superbubble (and not a superbubble).

#### 4.1 Missing proofs

**Lemma 1.** *Let $G$ be a bidirected graph, let $u \in V(G)$ be a vertex, and let $G'$ be the graph obtained from $G$ by flipping $u$. Let $W$ be a sequence of edges in $G$ and let $W'$ be the same sequence of edges in $G'$. Then $W$ is a walk in $G$ if and only if $W'$ is a walk in $G'$.*

*Proof.* Suppose $W = \{v_1\alpha_1, v_2\alpha_2\}, \ldots, \{v_{k-1}\hat{\alpha}_{k-1}, v_k\alpha_k\}$ is a walk in $G$. Whenever $v_i = u$ we have that $u$ is entered with sign $\hat{\alpha_i}$ and left with sign $\alpha_i$ in $G'$ because flipping $u$ swaps its positive vertex-sides with its negative vertex-sides, and whenever $v_i \neq u$ we have that $v_i$ is entered with sign $\alpha_i$ and left with sign $\hat{\alpha_i}$ in $G'$, and thus $W'$ is a walk in $G'$. By an identical argument, if $W'$ is a walk in $G'$ then $W$ is a walk in $G$. □

**Proposition 1.** *Let $G$ be a bidirected graph having at least two vertices. If $G$ has at most one tip then $G$ has a cycloid.*

*Proof.* Suppose that $x$ is a tip in $G$. Let $p$ be a maximal path in $G$ starting in $x$ (i.e., a path that cannot be made longer). This path has at least two vertices because $G$ has at least two vertices (and is connected, as we always assume implicitly). Further, the last vertex on this path, say $y$, is not a tip since $x$ is the unique tip in $G$. Suppose that $y$ is entered in $p$ with a vertex-side $y\beta$. Then every edge $\{y\hat{\beta}, z\gamma\}$ of $G$ is such that $z$ is in $p$, for otherwise $p$ is not maximal. But extending $p$ with such an edge results in a cycloid, where the possible exception occurs in $z$. If $G$ has no tip then we can pick an arbitrary vertex as the first vertex of a maximal path and proceed identically as above. □

**Lemma 2.** *Algorithm 1 only changes $G^\star$ by flipping vertices or by adding tips via subdivision of edges.*

*Proof.* Follows by examining Algorithm 1. □

**Lemma 3.** *At the end of Algorithm 1, every edge of $G^\star$ has opposite signs in its vertex-sides, that is, every edge of the input graph becomes directed.*

*Proof.* Consider an edge $e = \{u\alpha, v\beta\}$ when it is processed in ORIENTDFS. Notice that a vertex is flipped at most once during the algorithm since the DFS only flips vertices at the time of their first visit (i.e., when flipped[·] is null).

Suppose $v$ is unvisited. If $\alpha = \beta$ then the algorithm flips $u$, thus making $e$ a directed edge in $G^\star$. Otherwise $\alpha \neq \beta$ meaning that $e$ is already directed.

Suppose $v$ is visited. Then $v$ cannot be flipped. If $\alpha = \beta$ then $e$ is removed from $G^\star$ and in turn two directed edges are added: $\{u+, w-\}$ and $\{v+, w-\}$ if $\alpha = +$, and $\{u-, w+\}$ and $\{v-, w+\}$ otherwise. Otherwise $\alpha \neq \beta$ and so $e$ is directed.

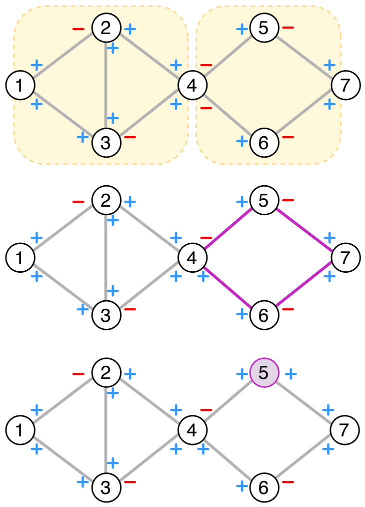


Figure 5: **Illustration of Lemma 4.** Assume for a contradiction that $\{1+, 7+\}$ is an ultrabubble and vertex 4 is a cutvertex of its ultrabubble component $X$. If like on top, the signs incident to 4 are partitioned according to the cut, then minimality contradicts $\{1+, 7+\}$ being an ultrabubble. Otherwise, if the signs are not partitioned according to the cut, either $X$ has a cycloid like the violet walk in the middle, or $X$ has a tip like the violet vertex 5 on the bottom, both contradicting $\{1+, 7+\}$ being an ultrabubble.

All cases have been examined. Notice that every bidirected edge of $G$ processed by the DFS either is already directed, becomes directed due to flipping, or is exchanged by two directed edges. Since every edge of $G$ is processed during the traversal, every edge of $G^\star$ is directed at the end of the search. □

**Lemma 4.** *Let $G$ be a bidirected graph and let $\{u\alpha, v\beta\}$ be pair of vertex-sides satisfying conditions (a), (b), and (c) of ultrabubbles with $u \neq v$. Let $X$ denote the component of $\{u\alpha, v\beta\}$. Then $\{u\alpha, v\beta\}$ is an ultrabubble if and only if no vertex in the interior of $X$ is a $u$-$v$ cutvertex in $X$.*

*Proof.* We can assume that $X$ has at least three vertices, since the statement clearly holds otherwise.

($\Rightarrow$) Suppose for a contradiction that $x \in V(X)$ is a $u$-$v$ cutvertex in $X$. Notice that $x$ is not a tip since the interior of ultrabubbles do not contain tips. We claim that the graph $X - x$ has exactly two components, $X_u$ containing $u$, and $X_v$ containing $v$. Suppose otherwise and let $Y'$ be a third component. Let $Y$ denote the graph where we put back $x$ in $Y'$ as well as all edges incident to $x$ from the vertices in $Y'$, i.e., $Y := X[V(Y') + x]$. No vertex in $Y$ but possibly $x$ is a tip in $Y$, since otherwise this vertex is also a tip in $X$, a contradiction to the fact that the interior of ultrabubbles do not contain tips. So $Y$ has at most one tip and thus it has a cycloid by Proposition 1, a contradiction to the acyclicity of ultrabubbles since $Y \subseteq X$.

Let us redefine $X_u$ so that it contains $x$ and all edges incident to $x$ from vertices in $X_u$, and redefine $X_v$ analogously. See Figure 5 for an example of the cases that follow.

If $x$ is a tip in $X_u$ and $v$ is a tip in $X_v$, then, for some $\gamma \in \{+, -\}$, $x$ is a tip with sign $\gamma$ in $X_u$ and $v$ is a tip with sign $\hat{\gamma}$ in $X_v$ since $x$ is not a tip. So $\{u\alpha, x\gamma\}$ and $\{x\hat{\gamma}, v\beta\}$ are clearly separable and hence $\{u\alpha, v\beta\}$ is not minimal, a contradiction.

Otherwise, and without loss of generality, $x$ is not a tip in $X_v$. Moreover, $X_v$ has no tips except for $v$ because ultrabubble components do not contain tips in their interior. Applying Proposition 1 to $X_v$ gives a contradiction to the acyclicity of $X$ since $X_v \subseteq X$.

($\Leftarrow$) Suppose for a contradiction that $X$ has vertex-sides $z\gamma$ and $z\hat{\gamma}$ such that $\{u\alpha, z\gamma\}$ and $\{z\hat{\gamma}, v\beta\}$ are separable, say with components $X_\gamma$ and $X_{\hat{\gamma}}$, respectively. Notice that any $u\alpha$-$v\beta$ bidirected path $p$ in $X$ through $z$ enter $z$ with sign $\gamma$ and leaves it with sign $\hat{\gamma}$, for otherwise $X_\gamma$ or $X_{\hat{\gamma}}$ have at most one tip and thus have a cycloid by Proposition 1, a contradiction to the acyclicity of $X$. So the suffix of this bidirected path starting at $z$ is a $z\hat{\gamma}$-$v\beta$ bidirected path avoiding vertex $u$, and hence it avoids the vertex-side $u\hat{\alpha}$ and obviously the vertex side $z\gamma$ (in the rest of the argument it is enough to consider this bidirected path as an undirected path). Since $X$ has two internally-vertex disjoint $u$-$v$ undirected paths, $X$ has a $u\alpha$-$v\beta$

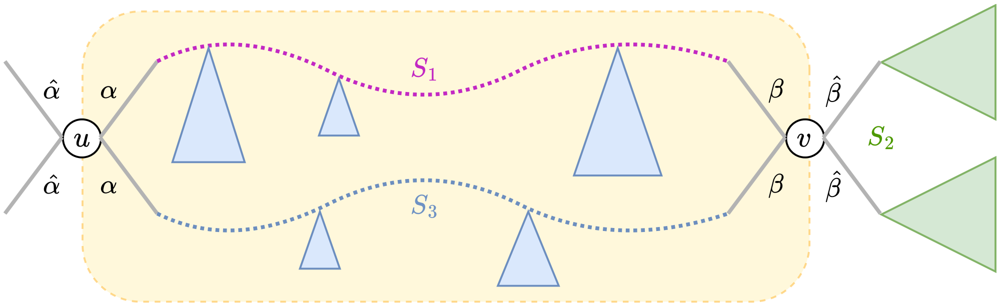


Figure 6: **Illustration of Lemma 5.**The notation $S_i$ for $i \in \{1, 2, 3\}$ denotes the segment of execution of the DFS from the point when it enters the ultrabubble (depicted in yellow). The segment $S_1$, in pink, corresponds to the first branch of the DFS tree. The segment $S_2$, in green, corresponds to the segment where the DFS exits the ultrabubble and traverses the rest of the graph (the green trees do not intersect the yellow region). The segment $S_3$, in blue, corresponds to the part where the DFS reenters the ultrabubble and visits every vertex and edge missed by $S_1$.

undirected path avoiding vertex $z$. So this path avoids the vertex-side $z\gamma$ and obviously the vertex-side $u\hat{\alpha}$. So $X \subseteq G$ has a $u\alpha$-$z\hat{\gamma}$ undirected walk avoiding $z\gamma$ and $u\hat{\alpha}$, and thus splitting $u\alpha$ and $z\gamma$ in $G$ leaves $z$ and $z'$ connected, a contradiction to the separability of $\{u\alpha, z\gamma\}$. □

**Corollary 1.** *The underlying undirected graph of an ultrabubble component is biconnected.*

*Proof.* Let $\{u\alpha, v\beta\}$ be an ultrabubble with component $X$. The statement is clear for trivial ultrabubbles, so suppose $V(X) \geq 3$.

Suppose for a contradiction that there is a vertex $x \in V(X) \setminus \{u, v\}$ such that $X - x$ is disconnected with components $C'_1, \ldots, C'_\ell$ ($\ell \geq 2$). Notice that $u$ and $v$ are not in different components of $X - x$ because $X$ has two internally vertex-disjoint $u$-$v$ undirected paths by Lemma 4. Let $C_i := X[V(C'_i) + x]$ for each $i \in \{1, \ldots, \ell\}$. Any component $C_i$ but the one containing $u$ and $v$ contains at most one tip (which is $x$) since ultrabubble components are tipless, and so it has a cycloid by Proposition 1. Since $C_i \subseteq X$, $X$ has a cycloid, a contradiction to the acyclicity of ultrabubble components. □

Given that ultrabubbles do not have cycloids with exception and contain exactly two tips, it is not hard to see that the following result holds (see also (Harary, 1953)).

**Proposition 2.** *Let $G$ be a bidirected graph with tips $u$ and $v$ of signs $\alpha$ and $\beta$, respectively. If $\{u\alpha, v\beta\}$ is an ultrabubble with component $G$ then Algorithm 1 does not subdivide any edge of $G$.*

*Proof.* Consider the first edge $e = \{x\gamma, y\gamma\}$ that the algorithm subdivides, and let $E'$ be the union of $\{e\}$ and the set of edges that the algorithm has already oriented, that is, we have already fixed whether we flip the endpoints of the edge and the vertex-sides have opposing signs. Note that the subgraph $G'$ of $G$ edge-induced by $E'$ has no tips except $u$ and $v$ due to the algorithm always considering the edges of the opposite sign first.

Consider now any maximal bidirected paths $P_x$ and $P_y$ in $G'$ starting from $x$ with vertex-side $x\hat{\gamma}$ and from $y$ with vertex-side $y\hat{\gamma}$, respectively. (If these were to repeat a vertex, they would produce a cycloid, a contradiction to acyclicity.) Since only $u$ and $v$ are tips, both walks end at $u$ or $v$. If both end at the same vertex, then the concatenation of $P_x$, $\{e\}$, and $P_y$ forms a cycloid, a contradiction. Therefore, one walk ends at $u$ and the other at $v$. Because all the other edges than $e$ in $E'$ have different signs for the two vertex-sides, the walks reach $u$ and $v$ with vertex-sides $u\gamma$ and $v\gamma$.

This, however, results in a contradiction, since $E'$ contains a $u\gamma$-$v\hat{\gamma}$ bidirected walk or a $u\hat{\gamma}$-$v\gamma$ bidirected walk (and thus there can be no same sign $\gamma$ incident to both $u$ and $v$ in $G'$ since they are both tips). To see this, consider the following. Without loss of generality, we can assume the algorithm starts the depth-first search from the tip $u$. Until the recursion reaches $v$, the first time the algorithm exits its current vertex $w$, it exits $w$ with an opposite sign than with which $w$ was entered and further goes to an unvisited vertex, since otherwise $w$ would be a tip or $G$ would contain a cycloid. In other words, the depth-first search first finds a $u\alpha$-$v\hat{\alpha}$ bidirected walk, where the latter sign follows from the algorithm flipping some of the vertices on the walk. Notably, no subdivision occurs before this walk is found otherwise there is a cycloid, and by construction one of the tips has a sign different from $\gamma$, possibly due to flipping the vertices. □

**Lemma 5.** *Let $G$ be a bidirected graph and let $\{u\alpha, v\beta\}$ be an ultrabubble with ultrabubble component $X$. Then Algorithm 1 does not subdivide any edge of $X$.*

*Proof.* Notice that whenever the DFS visits for the first time a vertex $w \neq r$ through a vertex-side $w\hat{\gamma}$ with $w\gamma$ being a vertex-side of an ultrabubble then the next vertex stacked by the DFS is in $N_G^{\gamma}(w)$,[1] so it is in fact a vertex contained in the interior of the ultrabubble. For the case $w = r$, suppose that $r$ is a tip with sign $\gamma$ and that $r\gamma$ is a vertex-side of an ultrabubble. If $\gamma = -$ then the claim also holds because the DFS starts at $r+$ by construction, and if $\gamma = +$ then the the next vertex visited by the DFS is also in the interior of the ultrabubble since there are no vertices in $N_G^{-}(r)$ to be visited. (Intuitively, the DFS always enters ultrabubble components from the "outside".)

From the discussion above, we can consider the first time the DFS stacks the opposite vertex-side of the vertex-sides of the ultrabubble, say $u\hat{\alpha}$ without loss of generality (possibly $u\alpha = r\gamma$ although $r-$ is never stacked as discussed previously; and if $G$ has no vertex-side $u\hat{\alpha}$ then it is not hard to see that $u\alpha = r\gamma$ and the ultrabubble component of $\{u\alpha, v\beta\}$ is $G$, and that there are no tips in $G$ besides $v$ and $u = r$). So the DFS enters $X$ by processing a vertex in $N_G^{\alpha}(u) \subseteq V(X)$. Eventually $v\beta$ is stacked and by the separability of $\{u\alpha, v\beta\}$, the time between $u\alpha$ and $v\beta$ are stacked no vertex in $V(G) \setminus V(X)$ is processed (indeed, $v$ is always present in the first branch of $T$); call this segment of execution $S_1$.

The next vertex processed by the DFS is in $N_G^{\hat{\beta}}(v)$ and thus it is not vertex of $X$, i.e., the DFS exits $X$. Later, once every vertex in $N_G^{\hat{\beta}}(v)$ is unstacked, $v$ becomes active again and the DFS reenters $X$ by considering the vertices in $N_G^{\beta}(v)$; call $S_2$ the segment of execution between the point where the DFS exits and reenters $X$. Importantly, notice that during this time no vertex of $X$ is processed due to the separability of $\{u\alpha, v\beta\}$ and the fact that $u$ is still in the stack.

So now the vertices $w \in N_G^{\beta}(v) \subseteq V(X)$ are processed. Notice that, similarly to before, the separability of $\{u\alpha, v\beta\}$ and the fact that $u$ is still in the stack implies that the subtree rooted at each $w$ is contained in $X$. Thus, eventually each $w$ is unstacked and next so is $v$, with no vertex in $V(G) \setminus V(X)$ being unstacked in the middle. Hence the next vertex to be active is a $\beta$-neighbor of $v$ and so it is a vertex of $X$. Once this vertex is unstacked, another vertex of $X$ is processed and so on, until $u$ is active again, and then unstacked. Thus, at this point, every vertex of $X$ has been processed. Call $S_3$ the segment of execution between the point where the DFS reenters $X$ and unstacks $u$ (see Figure 6).

Notice that $S_2$ is completely independent of $S_1$ and $S_3$, and so we can consider the execution of Algorithm 1 on $G$ as if when entering $X$ for the first time it processes the whole graph $X$ at once, i.e., when the algorithm processes $u$ it executes $S_1$ followed directly by $S_3$ and only then executes $S_2$. Thus, this hypothetical execution of Algorithm 1 is essentially the same with respect to the subdivision of edges as if the algorithm is executed on input $X$ alone. (Intuitively, the DFS-tree produced by this swap is identical as the one produced without swap, so whatever edges are subdivided in $S_1, S_2, S_3$ are also subdivided in $S_1, S_3, S_2$.) Executing Algorithm 1 on $X$ implies that no edge of $X$ is subdivided by Proposition 2, and therefore no edge of $X$ is subdivided when executing Algorithm 1 on $G$. □

**Corollary 2.** *Ultrabubble components are digraphic.*

*Proof.* Let $G$ be an ultrabubble component and consider the execution of Algorithm 1 on $G$. By Lemma 2 and Proposition 2, Algorithm 1 only changes $G$ by flipping vertices. Since every edge of $G^{\star}$ is directed by Lemma 3, $G^{\star}$ witnesses that $G$ is digraphic. □

**Theorem 1.** *Let $G$ be a bidirected graph with at least one tip. Algorithm 2 finds all and only the ultrabubbles of $G$ and runs in time $O(|V(G)| + |E(G)|)$.*

*Proof.* The correctness follows from Theorem 1 and Theorem 2. For the running time, notice that $D$ has size $\Theta(|V(G)| + |E(G)|)$ since the DFS subdivides each edge of $G$ with a tip at most once. Moreover, the

[1] Recall the definition of $E_v$ in Algorithm 1.

**Algorithm 2** Computing ultrabubbles

**Require:** A bidirected graph $G$ containing a tip
**Ensure:** The set of ultrabubbles of $G$

```
𝒰 ← ∅
Call Algorithm 1 (from Main Paper) on G to obtain D and the array flipped
ℬ ← FindWeakSuperbubbles(D)
for (u, v) ∈ ℬ do
    if {u, v} ∩ Z ≠ ∅ then
        continue
    end if
    α ← + if flipped[u] is false and − otherwise
    β ← − if flipped[v] is false and + otherwise
    𝒰 ← 𝒰 ∪ {uα, vβ}
end for
return 𝒰
```

DFS itself clearly runs in time $\Theta(|V(G)| + |E(G)|)$ (notice that every vertex is flipped at most once). Since weak superbubbles can be computed in linear time in the size of the input graph (see (Gärtner and Stadler, 2019)) and since the retrieval of the ultrabubbles takes constant time per weak superbubble (by querying the array flipped), Algorithm 2 runs in time $O(|V(G)| + |E(G)|)$. □

#### 4.2 Handling graphs with cutvertices

So far we assumed that the bidirected graph $G$ has at least one tip. In this section we show that the orientation algorithm works also on graphs without tips, as long as there is a cutvertex. The idea is to start the orientation algorithm from a cutvertex $v$, but with a properly chosen sign $\alpha$ (i.e., to call $\textsc{OrientDFS}(v, \alpha)$), as we discuss below (see Figure 7 for an illustration).

Let $C'_1, \ldots, C'_k$ be the components of the input bidirected graph obtained after removing $v$, and let $C_i$ be the bidirected subgraphs of $G$ induced by the vertices of $C_i$ together with $v$, i.e. $C_i = G[V(C'_i) + v]$. If a sign $\beta \in \{+, -\}$ is such that there are edges with vertex-side $v\beta$ in two subgraphs $C_i$ and $C_j$, for $i \neq j$, then $v\beta$ cannot be a defining vertex-side of any ultrabubble: the other vertex of the ultrabubble is not in one component between $C_i$ and $C_j$, and thus that component has at most one tip (which is $v$) and hence it has a cycloid. If edges with vertex-side $v\alpha$ (for some $\alpha \in \{+, -\}$) appear in only one $C_i$, then $v\alpha$ is the only candidate vertex-side of an ultrabubble and we can safely start the orientation algorithm as $\textsc{OrientDFS}(v, \hat{\alpha})$, so that at $v$ we first traverse the edges incident to $v$ with sign $\alpha$ (i.e., we enter the ultrabubble, if any, with $v\alpha$; see the proof of Lemma 5).

In the last case we have only two subgraphs $C_1$ and $C_2$, and each sign in $\{+, -\}$ at $v$ appears in exactly one of $C_1$ and $C_2$. In this case, starting the orientation with any sign, say $\textsc{OrientDFS}(v, +)$ is correct, because $v$ acts like a tip in each component, and they are visited separately by the DFS orientation algorithm.

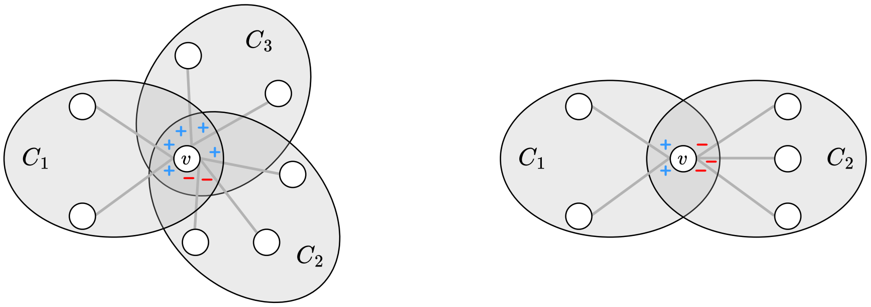


Figure 7: If a bidirected graph contains a cutvertex $v$, we can start the DFS orientation algorithm at $v$ in a special manner. In each illustration, we let $C_i$ be each of the subgraphs of the bidirected graph induced by each component of $G - v$, to which we add back vertex $v$ and the edges therein incident to it. On the left, we have the case where $+$ incident edges of $v$ appear in more than one component obtained by removing $v$. As such $v+$ cannot be the vertex-side of any ultrabubble. However, $-$ incident edges of $v$ appear only in one component, and thus $v-$ is a candidate ultrabubble vertex-side. We can start Algorithm 1 as OrientDFS$(v, +)$ so that we first traverse the $-$ edges incident to $v$. On the right hand side, each of $+$ and $-$ incident edges appear each in exactly one component obtained by removing $v$; starting the orientation algorithm with any sign is correct in this case ($v$ behaves like a tip in each component).

### 5 Graph statistics and bubble counts

We validated `BubbleFinder` by comparing its ultrabubble counts with `vg snarls` and between its orientation and doubled-graph modes (Table 1). On every dataset, the two `BubbleFinder` modes produced identical counts and ultrabubble sets, which also agreed with those reported by `vg snarls`. For HPRC v2.0, `vg snarls` did not finish on GFA input within 24 h, so the corresponding counts were obtained from GBZ input. BubbleGun, which reports nontrivial superbubbles, matches the nontrivial ultrabubble counts on most datasets. On four datasets, however, it reports one or two additional bubbles, which we traced to incorrect reports on graph structures that contain no ultrabubble. For example, in the HPRC v1.1 Chr. X graph, `BubbleGun` reports one additional bubble of type "super" with endpoints 204056176 and 204056248. To validate this, we extracted this bubble in a GFA file available at `https://github.com/algbio/BubbleFinder/blob/main/example/bubblegun-HPRC-v1.1-Chr.-X-bubble.gfa`. However, this is not an ultrabubble, since it contains bidirected edges $\{204056176+, 204056248-\}$ and $\{204056176-, 204056248-\}$, which make a cycloid. We isolated a similar problem even on a very small graph, which we show in Figure 8, and available as GFA file at `https://github.com/algbio/BubbleFinder/blob/main/example/bubblegun-small-bubble.gfa`.

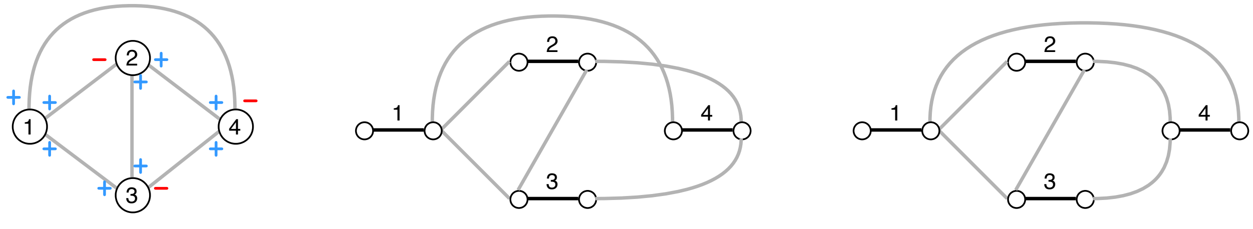


Figure 8: A graph without any ultrabubbles where BubbleGun v1.2.0 reports $\{1, 4\}$ as bubble of type "super". On the left we show the bidirected representation, on the middle the representation as a biedged graph (recall Figure 1), and on the right an equivalent drawing of the biedged graph.

Billi-heuristic, which reports nontrivial panbubbles, returns counts equal to or higher than the nontrivial ultrabubble counts, as expected because panbubbles are a superset of ultrabubbles. Topology statistics of the benchmark graphs are reported in Table 2.

Table 1: Number of bubbles reported by each tool. All counts are based on GFA input, except for HPRC v2.0 with `vg snarls`, which was run on GBZ input because the GFA run timed out after 24 h. Ultrabubble counts are shown both including and excluding trivial bubbles, as controlled by the `-T` flag in `vg snarls` and `BubbleFinder`. For `vg snarls`, the *snarls* and *diff* columns correspond to runs with trivial bubbles included; *diff* denotes the number of snarls that are not ultrabubbles. BubbleGun reports nontrivial superbubbles, and Billi-heuristic reports nontrivial panbubbles. TO = exceeded the time limit; N/A = tool does not run on graphs with tipless connected components. We mark in **bold** the values equal to the corresponding number of non-trivial ultrabubbles reported by `vg`.

| Type | Dataset | `vg snarls` | | | | `BubbleFinder` (ultrabubbles) | | | | `BubbleGun` | `Billi`-heuristic |
|---|---|---|---|---|---|---|---|---|---|---|---|
| | | ultrabubbles incl. trivial | snarls incl. trivial | diff | ultrabubbles excl. trivial | orientation incl. triv. | orientation excl. triv. | doubled incl. triv. | doubled excl. triv. | superbubbles | panbubbles |
| MC | HPRC v1.1 Chr. X | 1,692,356 | 1,692,401 | 45 | **1,623,552** | 1,692,356 | **1,623,552** | 1,692,356 | **1,623,552** | 1,623,553 | 1,623,592 |
| | HPRC v1.1 Chr. Y | 457,568 | 457,581 | 13 | **372,707** | 457,568 | **372,707** | 457,568 | **372,707** | **372,707** | 372,720 |
| | HPRC v1.1 CHM13 (47 indiv.) | 30,642,064 | 30,642,640 | 576 | **29,322,185** | 30,642,064 | **29,322,185** | 30,642,064 | **29,322,185** | **29,322,185** | 29,322,731 |
| | HPRC v2.0 CHM13 (232 indiv.) | 48,813,851 | 48,814,258 | 407 | **46,260,413** | 48,813,851 | **46,260,413** | 48,813,851 | **46,260,413** | TO (24 h) | 46,260,770 |
| pggb | E. coli (50 indiv.) | 469,226 | 469,893 | 667 | **469,074** | 469,226 | **469,074** | 469,226 | **469,074** | **469,074** | 469,739 |
| | Mouse Chr. 19 (17 indiv.) | 1,997,688 | 2,000,595 | 2,907 | **1,997,595** | 1,997,688 | **1,997,595** | 1,997,688 | **1,997,595** | **1,997,595** | 2,000,499 |
| | Primate Chr. 6 (14 indiv.) | 10,597,474 | 10,600,010 | 2,536 | **10,597,445** | 10,597,474 | **10,597,445** | 10,597,474 | **10,597,445** | **10,597,445** | 10,599,914 |
| | Tomato Chr. 2 (23 indiv.) | 760,258 | 760,962 | 704 | **760,186** | 760,258 | **760,186** | 760,258 | **760,186** | **760,186** | 760,865 |
| dbg | M. xanthus (10 indiv.) | 103,391 | 156,687 | 53,296 | **41,777** | 103,391 | **41,777** | 103,391 | **41,777** | **41,777** | 41,780 |
| vg | 1000GP Chr. 1 | 6,095,110 | 6,095,110 | 0 | **6,095,099** | 6,095,110 | **6,095,099** | 6,095,110 | **6,095,099** | **6,095,099** | **6,095,099** |
| | 1000GP Chr. 10 | 3,747,617 | 3,747,617 | 0 | **3,747,616** | 3,747,617 | **3,747,616** | 3,747,617 | **3,747,616** | **3,747,616** | **3,747,616** |
| | 1000GP Chr. 22 | 1,027,822 | 1,027,822 | 0 | **1,027,814** | 1,027,822 | **1,027,814** | 1,027,822 | **1,027,814** | **1,027,814** | **1,027,814** |
| Pangene | E. coli v1.1 | 7,237 | 7,259 | 22 | **400** | 7,237 | **400** | 7,237 | **400** | **400** | 403 |
| | M. tb 152m p0 | 3,804 | 3,822 | 18 | **133** | 3,804 | **133** | 3,804 | **133** | **133** | 151 |
| | M. tb 152m p1 | 3,908 | 3,920 | 12 | **84** | 3,908 | **84** | 3,908 | **84** | **84** | 93 |
| | M. tb 152p | 3,744 | 3,760 | 16 | **131** | 3,744 | **131** | 3,744 | **131** | **131** | 147 |
| | Human 100 | 17,974 | 18,198 | 224 | **90** | 17,974 | **90** | 17,974 | **90** | 92 | N/A |
| | Human 100+10% | 17,129 | 17,379 | 250 | **169** | 17,129 | **169** | 17,129 | **169** | 170 | 403 |
| | Human 472 | 17,956 | 18,180 | 224 | **85** | 17,956 | **85** | 17,956 | **85** | 86 | 270 |
| | Human 472+10% | 17,280 | 17,527 | 247 | **160** | 17,280 | **160** | 17,280 | **160** | **160** | 383 |

### References


S. G. Bhat, D. Mahajan, and C. Jain. Billi: Provably accurate and scalable bubble detection in pangenome graphs. *bioRxiv*, 2025. URL `https://doi.org/10.1101/2025.11.21.689636`.

F. Dabbaghie, J. Ebler, and T. Marschall. BubbleGun: enumerating bubbles and superbubbles in genome graphs. *Bioinformatics*, 38(17):4217–4219, 07 2022.

F. Gärtner and P. F. Stadler. Direct superbubble detection. *Algorithms*, 12(4):81, 2019.

F. Harary. On the notion of balance of a signed graph. *Michigan Mathematical Journal*, 2(2):143–146, 1953.

H. Li, M. Marin, and M. R. Farhat. Exploring gene content with pangene graphs. *Bioinformatics*, 40(7): btae456, 2024.

B. Paten, J. M. Eizenga, Y. M. Rosen, A. M. Novak, E. Garrison, and G. Hickey. Superbubbles, ultrabubbles, and cacti. *Journal of Computational Biology*, 25(7):649–663, 2018.

Table 2: Graph topology statistics. $n$ =number of segments (nodes), $m$=number of links (edges). Tips and cut vertices are counted on the bidirected graph. A connected component (CC) without tips has no degree-1 node. “without cut vertices” has no node whose removal disconnects the CC. Conflict vertices are those introduced by the orientation algorithm (Algorithm 1 in the main paper).

| Type | Dataset | $n$ | $m$ | # CCs | # tips | # cut vertices | CCs w/o tips | CCs w/o cut vtx | # conflict vertices |
|---|---|---|---|---|---|---|---|---|---|
| MC | HPRC v1.1 Chr. X | 5,124,974 | 7,100,260 | 1 | 2 | 1,396,038 | 0 | 0 | 70 |
| | HPRC v1.1 Chr. Y | 1,675,938 | 2,308,100 | 1 | 2 | 163,552 | 0 | 0 | 23 |
| | HPRC v1.1 CHM13 (47 indiv.) | 92,879,580 | 128,165,765 | 25 | 50 | 23,280,102 | 0 | 0 | 2,691 |
| | HPRC v2.0 CHM13 (232 indiv.) | 148,283,410 | 206,031,684 | 25 | 50 | 35,870,507 | 0 | 0 | 244,265 |
| pggb | E. coli (50 indiv.) | 1,560,532 | 2,117,858 | 1 | 13 | 289 | 0 | 0 | 4,312 |
| | Mouse Chr. 19 (17 indiv.) | 6,257,525 | 8,667,808 | 4 | 28 | 667 | 0 | 0 | 6,593 |
| | Primate Chr. 6 (14 indiv.) | 34,386,688 | 46,883,806 | 9 | 67 | 3,221 | 0 | 0 | 19,813 |
| | Tomato Chr. 2 (23 indiv.) | 2,331,264 | 3,188,060 | 1 | 21 | 67,100 | 0 | 0 | 5,453 |
| dbg | M. xanthus (10 indiv.) | 1,605,531 | 2,122,751 | 1 | 315,547 | 305,845 | 0 | 0 | 6,665 |
| vg | 1000GP Chr. 1 | 12,499,232 | 12,499,231 | 1 | 2 | 6,088,506 | 0 | 0 | 0 |
| | 1000GP Chr. 10 | 7,689,606 | 7,689,605 | 1 | 2 | 3,743,190 | 0 | 0 | 0 |
| | 1000GP Chr. 22 | 2,131,985 | 2,131,984 | 1 | 2 | 1,026,304 | 0 | 0 | 0 |
| Pangene | E. coli v1.1 | 13,003 | 19,433 | 22 | 61 | 148 | 0 | 1 | 2,328 |
| | M. tb 152m p0 | 4,272 | 4,724 | 1 | 2 | 138 | 0 | 0 | 80 |
| | M. tb 152m p1 | 4,195 | 4,625 | 3 | 5 | 178 | 0 | 0 | 49 |
| | M. tb 152p | 4,216 | 4,653 | 1 | 2 | 120 | 0 | 0 | 86 |
| | Human 100 | 19,091 | 20,883 | 3 | 33 | 9,387 | 1 | 0 | 645 |
| | Human 100+10% | 19,041 | 22,527 | 1 | 25 | 2,295 | 0 | 0 | 1,300 |
| | Human 472 | 19,055 | 41,729 | 5 | 33 | 10,021 | 0 | 0 | 669 |
| | Human 472+10% | 19,019 | 23,571 | 2 | 26 | 3,222 | 0 | 0 | 1,094 |
